# Neocortical long-range inhibition promotes cortical synchrony and sleep

**DOI:** 10.1101/2024.06.20.599756

**Authors:** Jacob M Ratliff, Geoffrey Terral, Arenski Vazquez Lechuga, Stefano Lutzu, Arena Manning, Nelson Perez Catalan, Gabriela Neubert da Silva, Soyoun Kim, Julie Mota, Matt Mallory, Bianca Stith, Charu Ramakrishnan, Gianna Mattessich, Lief E Fenno, Tanya Daigle, David A Stafford, Hongkui Zeng, Bosiljka Tasic, Staci Sorensen, Karl Deisseroth, John Ngai, Thomas S. Kilduff, Lucas Sjulson, Stephanie Rudolph, Renata Batista-Brito

**Affiliations:** Albert Einstein College of Medicine, New York City, NY, United States; Biosciences Division, SRI International, Menlo Park, CA 94025, United States; Stanford University, Stanford, United States; The University of Texas at Austin, Austin, TX, United States; Allen Institute for Brain Science, Seattle, WA, United States; National Institute of Neurological Disorders and Stroke, National Institutes of Health, Bethesda, MD, United States; University of California, Berkeley, Berkeley, CA United States

**Author notes:** UCL Queen Square Institute of Neurology, University College London, London, UK. These authors are listed as co-first (Jacob M Ratliff, Geoffrey Terral). Corresponding author Correspondence should be addressed to: Renata Batista-Brito, Geoffrey Terral.

## Abstract

Sleep and wakefulness are associated with distinct cortical patterns of rhythmic activity. During low-arousal states such as slow wave sleep, synchronous low-frequency rhythms dominate activity across widespread cortical regions. Although inhibitory neurons are increasingly recognized as key players of cortical state, the *in vivo* circuit mechanisms coordinating synchronized activity across local and distant neocortical networks remain poorly understood. Here, we show that somatostatin and chondrolectin co-expressing cells (Sst-Chodl), a sparse and genetically distinct class of neocortical GABAergic inhibitory neurons, are selectively active during low-arousal states and largely silent during periods of high arousal. In contrast to most neocortical inhibitory neurons, Sst-Chodl cells, despite being extremely sparse, exert widespread influence across the neocortex via long-range axons that simultaneously target multiple regions. Selective activation of Sst-Chodl cells is sufficient to promote multi-region cortical synchronization characteristic of low-arousal states and to induce sleep. Together, these findings show that long-range Sst-Chodl inhibitory neurons not only track behavioral state but can actively promote sleep-like cortical activity and sleep behavior, highlighting an important contribution of cortical circuits to sleep regulation alongside established subcortical mechanisms.

## Introduction

Mammals spend much of their day in sleep and other states of rest. Regular engagement in restful states is essential for metabolism, hormone regulation, learning, positive emotion, attention, immune health, and other essential functions ^1–6^. Disruption of sleep and rest is associated with impaired physiological and cognitive function and is linked to a wide range of disorders, including insomnia, sleep apnea, narcolepsy, depression, ADHD, schizophrenia, and even death ^6–15^. Transitions between alertness and low-arousal states, such as quiet wakefulness and sleep, occur on the timescale of seconds and are accompanied by pronounced changes in neocortical activity patterns (i.e., oscillatory activity or cortical states) ^16–19^. Low-arousal states, including slow wave sleep (SWS) and quiet immobility, are characterized by synchronous low-frequency cortical fluctuations and coordinated spiking activity, often referred to as synchronized cortical states. These alternate with desynchronized states associated with periods of high arousal and active behavior, during which low-frequency cortical activity is suppressed ^16,17^. GABAergic inhibitory neurons (INs) have been implicated as key regulators of arousal-dependent neocortical activity ^20–25^. In particular, fast-spiking parvalbumin-expressing INs have been shown to promote desynchronized states by enhancing gamma-band activity ^26,27^. In contrast, far less is known about the identity and function of IN populations that regulate synchronized cortical states ^25^. Thus, the circuit mechanisms coordinating low-arousal, synchronized activity across local and distributed neocortical networks remain poorly understood. Here, we identify a small and distinct class of neocortical INs that is selectively active during low-arousal, synchronized states and whose activation is sufficient to promote synchronous neocortical activity and sleep.

Although the diversity of cortical INs has been recognized for nearly a century ^28^, assigning specific functions to individual IN classes has proven challenging, in part because established IN classes are themselves highly heterogeneous ^29–31^. For example, somatostatin (Sst)-expressing INs, which are often treated as a monolithic group, can be subdivided into >10 subclasses with distinct morpho-electric and transcriptomic properties ^24,29,30,32–34^. In this study, we focus on a transcriptionally homogeneous and evolutionarily conserved subtype of Sst INs, characterized by the co-expression of Sst, chondrolectin (Chodl), neuronal nitric oxide synthase (Nos1), and the Neurokinin-1 receptor (Tacr1); hereinafter here referred to as Sst-Chodl cells, corresponding to the NOS1-expressing cortical IN population described in previous studies ^30,32,33^. Sst-Chodl cells are GABAergic INs that differ strikingly from canonical, locally-projecting neocortical interneurons in that they possess long-range projection axons that can extend over millimeters in the mouse cortex and across centimeters in larger brains ^35–39^. Despite their extreme sparsity, comprising less than 1% of cortical GABAergic neurons, ^33,40^ and the substantial energetic cost associated with maintaining long-range projections, Sst-Chodl cells are highly conserved across vertebrate species, from salamanders to humans ^41,42^. Previous *ex vivo* studies have suggested that putative Sst-Chodl cells are likely active during SWS ^43–46^, raising the possibility that they contribute to synchronized states. However, their activity and functional roles *in vivo* have remained difficult to investigate due to the lack of tools for their specific targeting.

Here, we use an intersectional genetic strategy ^47–49^ to selectively target Sst-Chodl cells and determine their functional role within the neocortex. We show that Sst-Chodl cells form dense regional and inter-areal intracortical connections and are active during periods of high cortical synchrony, including quiet wakefulness and SWS. Activation of Sst-Chodl cells in the visual cortex drives both regional and large-scale neocortical network activity towards a sleep-like state characterized by enhanced cortical synchrony. Moreover, widespread activation of these cells across the neocortex promotes behavioral sleep. Together, these findings identify Sst-Chodl cells as a cortical circuit element well positioned to coordinate state-dependent network activity and associated behaviors. Consistent with a growing body of work indicating that cortical activity contributes to sleep regulation^50–58^, our results demonstrate that the activity of a specific subclass of neocortical INs, the Sst-Chodl cells, is sufficient to generate low-arousal cortical states and promote sleep.

## Results

### Sst-Chodl cells are long-range inhibitory neurons with dense projections

We used intersectional genetics to selectively target Sst-Chodl cells (Fig 1a, Supp Fig 1). Single-cell sequencing showed that labeling with the Nos1^creER^ mouse line captured a heterogeneous population in which Sst-Chodl cells represented only a minority, whereas the double-transgenic mouse lines combining Sst^flp^ with Chodl^cre^ or Nos1^creER^ achieved similarly high specificity for Sst-Chodl targeting (Fig 1a) ^49^. Immunohistochemistry confirmed robust and selective expression in Sst-Chodl cells (Supp Fig 1d–e).

**Figure 1.**
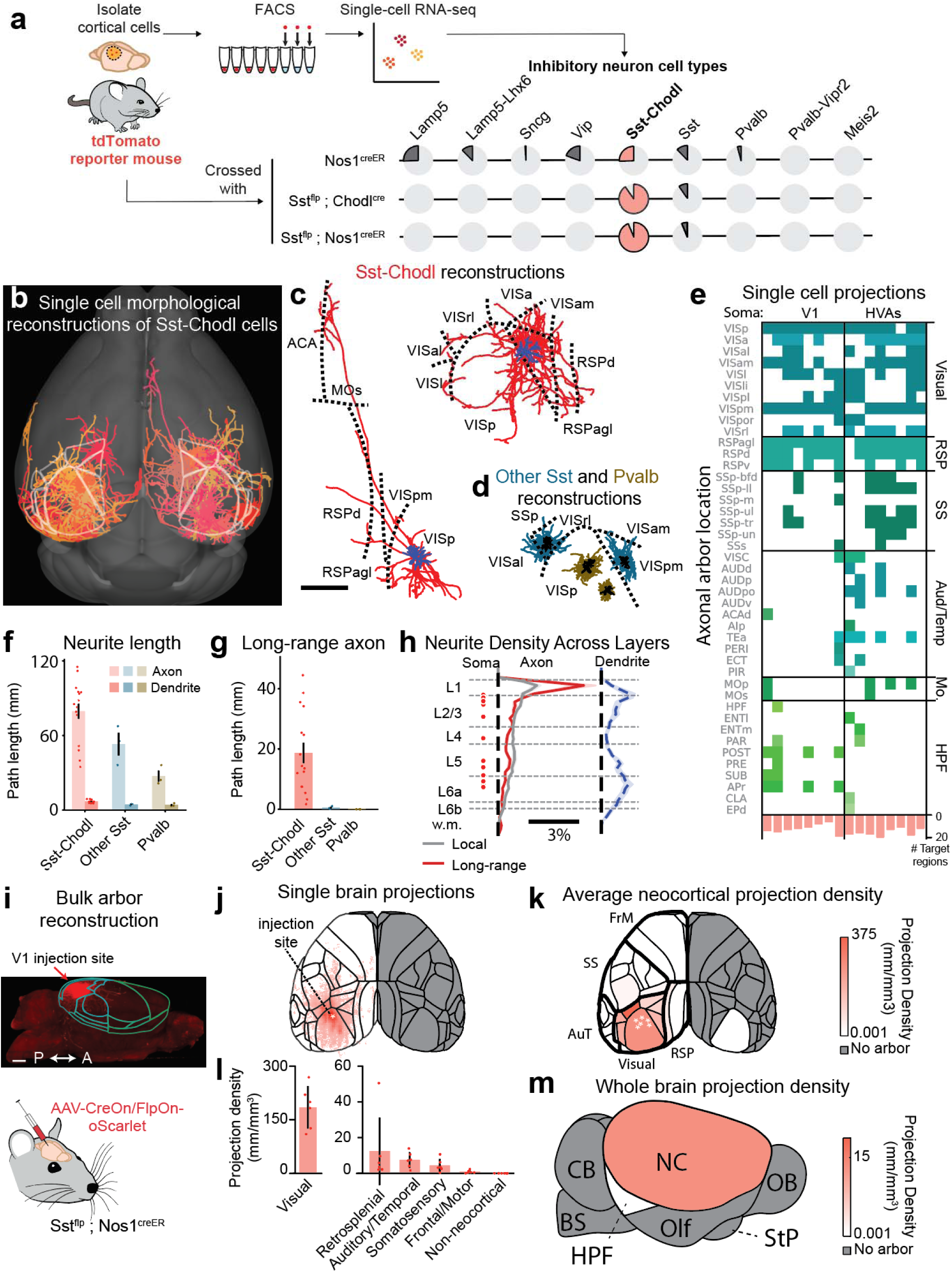
Sst-Chodl cells are long-range inhibitory neurons with extensive intracortical projections. **a)** Experimental strategy for transcriptomic profiles and quantification of inhibitory neuron cell types labeled from single cell RNA sequencing from tdTomato reporter mice crossed with Nos1^CreER^ or Sst^flp^ ; Chodl^cre^ or Sst^flp^ ; Nos1^creER^ mouse lines (data from ^90^). These approaches selectively and robustly label Sst-Chodl cells. **b)** Single-cell reconstructions of Sst-Chodl cells in primary visual cortex (V1) and higher visual areas (HVAs). **c)** Example reconstructions of two Sst-Chodl cells (axon in red, dendrite in dark blue; scale: 1mm) illustrating dense local arborization together with long-range projections spanning millimeters. Dashed lines indicate area borders. **d)** Reconstructions of other Sst neurons (non-Chodl) (blue, dendrite in black) and a Pvalb neurons (khaki; dendrite in black), which show predominantly local axons with minimal inter-areal spread. These comparisons reveal that Sst-Chodl cells, unlike other inhibitory types, routinely project far beyond their home cortical area. **e)** Axonal arborization matrix summarizing downstream cortical regions targeted by individual Sst-Chodl cells (columns); colored boxes indicate presence of axon in region. Bottom: number of downstream regions innervated by each reconstructed cell. **f)** Comparison of axon and dendrite length across Sst-Chodl, other Sst and Pvalb cells. **g)** Axon length located outside of visual cortical areas. **h)** Layer distribution of axons, dendrite density and soma positions for reconstructed neurons (N = 16 Sst-Chodl cells, 3 other Sst cells, and 3 Pvalb cells). These analyses show that Sst-Chodl cells have widespread intracortical arbors, with long-range axons traveling preferentially through superficial layers. **i)** Strategy for bulk arbor reconstruction following AAV-CreOn/FlpOn-oScarlet injection into V1 of Sst^flp^ ; Nos1^CreER^ mice, with example sagittal max-intensity projection of labeled fibers (scale 1mm, A: anterior, P: posterior). **j)** Thin-section reconstructions showing projection patterns in an example brain. **k)** Mean projection density across brain, with inferred injection sites marked (*). **l)** Quantification of projection densities within visual cortex and across other brain areas. **m)** Brain-wide distribution of V1 Sst-Chodl projections (N = 6). These population-level datasets confirm that Sst-Chodl cells send dense local axons within visual areas and widespread ipsilateral projections across most of the neocortex, with sparse extensions into para-hippocampal regions. Data are means; shading and bars indicate s.e.m. Abbreviations: From c): ACA, anterior cingulate area; Mos, secondary motor area; RSPd, retrosplenial area (dorsal); RSPagl, retrosplenial area (lateral agranular); SSp, primary somatosensory area; VIS, visual area (further divided into pm = posteromedial, p = primary, l = lateral, al = anterolateral, rl = rostrolateral, a = anterior, am = anteromedial). From e) Mo, motor cortical areas; SS, somatosensory cortical areas; Aud/Temp, auditory and temporal cortical areas; RSP, retrosplenial cortical areas; HPF, hippocampal formation. From k) FrM, frontomotor cortex; AuT, auditory and temporal cortical areas. From m): CB, cerebellum; NC, neocortex; OB, olfactory bulb; BS, brainstem; HPF, hippocampal formation; Olf, olfactory areas; StP, striatum/pallidum.

Next, to evaluate the extent of Sst-Chodl cell projections, we used the Sst^flp^ ; Chodl^cre^ intersection to sparsely label individual neurons, enabling single-cell anatomical reconstruction from fluorescent Micro-Optical Sectioning Tomography (fMOST) datasets annotated in HortaCloud (see Methods) ^59,60^. This approach allowed us to resolve the complete local and long-range axonal arbors of individual Sst-Chodl cells across the neocortex. We reconstructed the full morphology of 16 Sst-Chodl cells across all layers^32^ in primary visual cortex (V1) and higher visual areas (HVA, Fig 1b) and compared their projection pattern with parvalbumin (Pvalb) and non-Chodl Sst (other Sst cells) reconstructed cells (Fig 1c-h). While Pvalb and other Sst cells had local arborization, rarely crossing area boundaries, we found that Sst-Chodl cells had both dense local arborization within visual areas and long-range axons traveling millimeters from the soma across broad neocortical areas (Fig 1b-f, Supp Table 1-2). The projections of Sst-Chodl cells were exclusively intracortical and ipsilateral, broadly targeting nearly all cortical areas, with the notable exception of far anterior (e.g., frontal pole, prelimbic area) and ventral (e.g., gustatory area, Fig 1e) regions. Importantly, projection patterns were highly similar between Sst-Chodl cells originating from primary visual cortex (V1) and higher visual areas (HVAs) (Supp Fig 2a-b, Supp Table 1-2) or between medial and lateral regions (Supp Fig 2c-f, Supp Table 1-2). We encountered local/regional projections across all layers, whereas the long-range projections were preferentially found in upper layers (Fig 1h).

**Figure 2.**
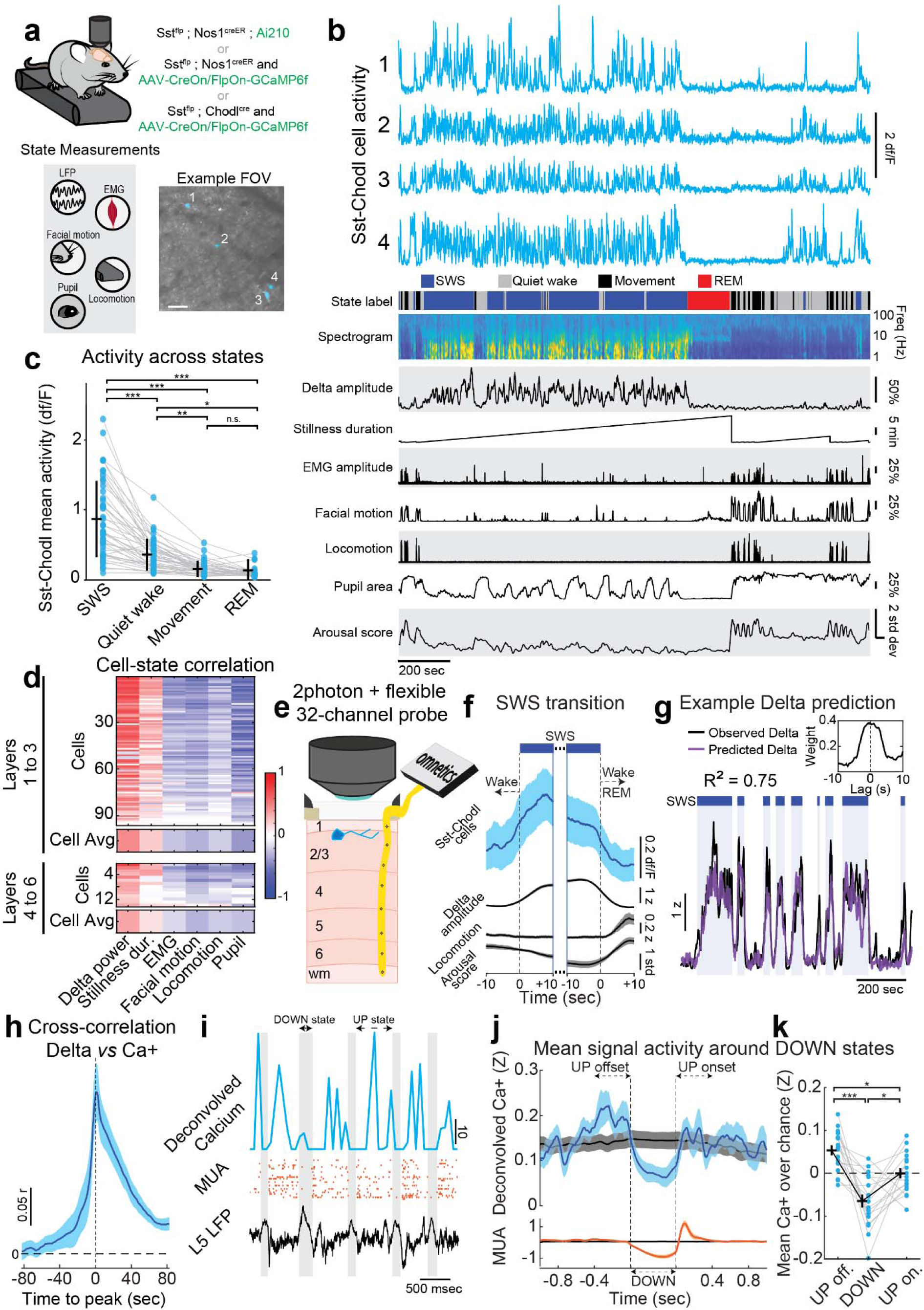
Sst-Chodl cells are strongly active during low-arousal, synchronized cortical states. **a)** Strategy for expressing GCaMP in Sst-Chodl cells and for simultaneous 2-photon imaging in V1, LFP recording in contralateral V1, and behavioral state monitoring in head-fixed animals. Example field of view (FOV) showing four labeled Sst-Chodl cells (ROIs in blue; scale: 100 µm). **b)** Example traces from the four imaged Sst-Chodl cells in (a), alongside behavioral/physiological measurements of sleep-wake states, which were combined into a one-dimensional arousal score. These measurements illustrate that Sst-Chodl activity increases when the cortex enters synchronous states such as SWS and Quiet Wake. **c)** Mean activity of layer 1-3 Sst-Chodl cells across behavioral states (ANOVA p < 0.001; post hoc comparisons: SWS-QW p < 0.001, SWS-Move p < 0.001, SWS-REM p < 0.001, QW-Move p = 0.001, QW-REM p = 0.010, Move-REM p = 0.990). This shows that Sst-Chodl cells in superficial layers are most active during SWS and Quiet Wake and suppressed during Movement and REM. **d)** Correlation between Sst-Chodl cell activity and state metrics (i.e., delta power, pupil diameter, locomotion, EMG) for cells in superficial layers (recorded through cranial windows, Layer 1-3) and in deeper cell layers (recorded through microprism, Layer 4-6). Sst-Chodl cell activity across all layers was positively correlated with markers of low arousal and negatively correlated with high-arousal signals. Layers 1-3: N = 15 animals (11 Sst^flp^ ; Nos1^creER^ mice, 4 Sst^flp^ ; Chodl^cre^ mice), n = 97 cells (76 Sst^flp^ ; Nos1^creER^, 21 Sst^flp^ ; Chodl^cre^); Layers 4-6: N = 5 animals (Sst^flp^ ; Chodl^cre^ mice), n = 14 cells. **e)** Schematic of simultaneous 2-photon imaging and LFP recording using a flexible 32-channel silicon probe inserted under the cranial window**. f)** Mean Sst-Chodl cell activity, delta power amplitude, locomotion, and arousal level aligned to transitions into and out of SWS. Sst-Chodl cell activity increases as animals enter SWS in parallel with rising delta power and decreases when they transition to desynchronized states. **g)** Cross-validated linear regression model predicting delta amplitude from deconvolved Sst-Chodl calcium signals, showing that activity reflects ongoing delta fluctuations rather than predicting future changes. **h)** Cross-correlation between delta amplitude and Sst-Chodl calcium activity, indicating tight temporal coupling. **i)** Example simultaneous recordings of Sst-Chodl cell calcium activity (deconvolved signal), multi-unit activity (MUA), and layer 5 LFP using flexible probes. DOWN states were detected from MUA and LFP signatures**. j)** Average peri-stimulus time histogram (PSTH) of calcium activity across Sst-Chodl cells (blue) and MUA (orange) aligned to DOWN state onset; chance levels shown in black. **k)** Average calcium activity from 400 msec before DOWN states (UP offset), during DOWN states, and 400 msec after DOWN states (UP onset). Sst-Chodl cells show increased activity prior to DOWN states (ANOVA p < 0.001; Bonferroni post hocs: UP off.-DOWN p < 0.001, DOWN-UP on. p = 0.017, UP off.-UP on. p = 0.036), consistent with engagement at the termination of UP states rather than during the silent DOWN phase. Panels f-k include data from 2-photon-probe recordings shown in in (e). N = 4 animals (Sst^flp^ ; Chodl^cre^ mice), n = 17 cells. ***: p < 0.001, **: p < 0.01, *: p < 0.05. Data are means; shading and bars indicate s.e.m.

After characterizing the single cell projection patterns, we next quantified the density of the projections from Sst-Chodl cells originating in V1 across the neocortex. To this end, we injected the intersectional AAV8-Ef1a-CreOn/FlpOn-oScarlet vector into V1 of Sst^flp^ and Nos1^creER^ double transgenic mice (hereafter “Sst^flp^ ; Nos1^creER^ mice”; Supp Fig 1), driving robust cytosolic labeling of a larger population of Sst-Chodl cells (Fig 1i). We annotated the oScarlet-labeled arbor of Sst-Chodl cells across the brain and confirmed that projections of V1 Sst-Chodl cells targeted broadly ipsilateral neocortical areas (with small contralateral V1 projections) and densely innervated visual areas (200-600 mm of arbor per mm^3^ of tissue; Fig 1j-l, Supp Fig 3). At the population level, Sst-Chodl cells exhibited extensive inter-areal projections that targeted most of the neocortex, including retrosplenial, auditory, somatosensory, and frontal motor areas (Fig 1l). Notably, only a limited set of far anterior and dorsal neocortical regions lacked detectable innervation (agranular insular area, frontal pole, gustatory areas, infralimbic area, orbital area). Consistent with the single cell reconstruction data (Fig 1b-g), we also observed small but reproducible projections from V1 Sst-Chodl cells beyond the neocortex, targeting ipsilateral para-hippocampal structures, including the subiculum and entorhinal cortex (Fig 1m, Supp Table 3).

Using whole brain tissue clearing and light sheet imaging, we compared the arborization patterns from animals with Sst-Chodl cells labeled in V1 and from animals with cells labeled in primary somatosensory cortex (S1) (Supp Fig 4, Supp Movie 1). Similarly to cells in V1, we found that Sst-Chodl cells in S1 had dense arbors spanning all layers within the injected regions and long-range interareal projections travelling preferentially through superficial layers (Supp Fig 4b-f). Those projections were ipsilateral and confined to the neocortex (Supp Fig 4b-g). While inter-areal projections from V1 Sst-Chodl cells extensively targeted retrosplenial areas, the long-range projections of S1 Sst-Chodl cells more densely innervated areas surrounding the injection site such as frontal, motor and visual areas (Supp Fig 4b-g).

Together, these data demonstrate that Sst-Chodl cells originating from distinct neocortical areas share a common feature: they are long-range projecting neurons that broadly target the ipsilateral neocortex.

### Sst-Chodl cells receive neocortical inputs from diverse cell classes

To identify the presynaptic partners of Sst-Chodl cells, we employed an intersectional genetic monosynaptic retrograde tracing strategy combined with whole-brain imaging and registration to the Allen Institute Common Coordinate Framework (Supp Fig 5a, Supp Table 4). Tracing was performed from Sst-Chodl cells in primary visual cortex (V1), enabling systematic mapping of their upstream inputs. Presynaptic neurons were almost exclusively located within neocortical areas, with only a small fraction detected in the thalamus (<4% of all labeled cells; Supp Fig 5b-c). Within the neocortex, labeled neurons were restricted to the ipsilateral hemisphere, distributed primarily across intermediate layers (2–5); and most densely represented in V1 (Supp Fig 5d–i). Immunohistochemical analyses revealed that presynaptic neurons comprised both glutamatergic (Ctip2⁺: putative pyramidal tract, Satb2⁺: putative intratelencephalic) and GABAergic (Pvalb⁺, Vip⁺, Sst⁺) populations, with the majority co-expressing Satb2 and Vip, and the fewest co-expressing Sst and Nos1 (Supp Fig 5j,k).

Together, these data indicate that Sst-Chodl cells integrate a broad diversity of regional excitatory and inhibitory inputs from all the major glutamatergic and GABAergic classes that we probed.

### Sst-Chodl cells are active during periods of low-arousal and high neocortical synchrony

To test whether Sst-Chodl cell activity varies across sleep and wake states, as suggested by previous c-Fos studies ^45,46^, we expressed a GCaMP calcium indicator using either viral vectors or transgenic line (see Methods) in Sst^flp^ ; Nos1^creER^ or Sst^flp^ ; Chodl^cre^ mice. We then used 2-photon calcium imaging to measure the activity of labeled cells across arousal and network synchronization states, measured using pupillometry, locomotion, contralateral cortical local field potentials (LFPs; delta band, 1–4 Hz), electromyography (EMG), and facial movements (Fig 2a-b, Supp Movie 2). Because these state measurements were strongly correlated (Supp Fig 6a), we combined them into a 1-dimensional “arousal score”, corresponding to the first principal component of these state variables (Fig 2b, Supp Fig 6b).

To facilitate sleep during imaging, mice were extensively habituated to head fixation and the recording setup (see Methods). Sleep states were scored automatically ^61,62^ and subsequent curated manually. Based on the combined state measurements, behavioral state was classified into four categories: wakefulness, subdivided into Movement and Quiet Wakefulness (Quiet Wake or QW); slow-wave sleep (SWS), and rapid eye movement (REM) sleep (Fig 2b). Wakefulness was characterized by high EMG activity and low delta power or shallow power spectral slope, with Movement distinguished by periods of high locomotion and facial movements, and QW defined by near-complete absence of motion. SWS was characterized by low EMG activity and high delta power or steep power spectral slope; whereas REM sleep was identified by low EMG activity and elevated theta power ^61,62^.

We found that a large proportion of labeled cells in superficial layers (L1-2/3) from both intersectional mouse lines exhibited intense activity during low-arousal, highly synchrony states such as SWS and Quiet Wake, and were strongly suppressed during REM sleep and Movement, when the neocortex is desynchronized (95 out of 111 cells; Fig 2b-c, Supp Fig 6c). A minor subset of labeled cells showed the opposite pattern of activation, being most active during periods of high movement (16 out of 111 cells; Supp Fig 6c-d). This subset is consistent with our sequencing and immunohistochemistry analyses indicating the presence of a minor population of non-Sst-Chodl cells in both intersectional crosses, as well as with a previously described small heterogenous populations of Sst⁺Nos1⁺ neurons that do not express Chodl ^32,63^ (Supp Fig 6e).

We next examined the consistency of Sst-Chodl cell activity across cortical layers using microprism-based imaging (Supp Fig 6f). Similar to superficial layer neurons, Sst-Chodl cells recorded in deep layers were strongly active during SWS and Quiet Wake and suppressed during Movement and REM sleep (14 out of 16 cells; Supp Fig 6g). Across all layers, Sst-Chodl cells active during low-arousal states exhibited a remarkably homogeneous activity profile, showing strong positive correlations with low-arousal metrics, including delta-band spectral power and time elapsed since last locomotion (i.e., stillness duration), and negative correlations with high-arousal measures such as EMG, facial motion, locomotion and pupil diameter (Fig 2d). Activity among these cells was also highly correlated and dependent on the animal’s arousal level, with stronger correlations observed during low-arousal states (Fig 2d, Supp. Fig 6h). Consistent with this relationship, separating LFP epochs based on periods of high versus low Sst-Chodl cell activity revealed that these neurons were most active when cortical networks were highly synchronized, as indicated by elevated delta power (i.e., high delta power; Supp Fig 6i-k).

To examine the temporal relationship between Sst-Chodl cell activity and cortical dynamics, we simultaneously recorded Sst-Chodl calcium signals and LFPs from the same cortical region (ipsilateral V1). Flexible 32-channel probes^64^ were implanted beneath the imaging window, enabling high-temporal resolution 2-photon imaging of Sst-Chodl cell activity alongside electrophysiological recordings (see Methods; Fig 2e-k). Sst-Chodl cells exhibited state-dependent changes in activity that closely tracked fluctuations in LFP in delta power (Fig 2f and Supp Fig 6l). As mice entered SWS, both Sst-Chodl cell activity and delta power increased and remained elevated throughout sleep bouts. Conversely, transitions into desynchronized states (wake or REM sleep) were accompanied by concurrent decreases in both Sst-Chodl cell activity and delta power (Fig 2f). Sst-Chodl cell activity closely co-varied with delta power rather than anticipating future changes in cortical synchrony. Consistent with this interpretation, a time-resolved generalized linear model showed that Sst-Chodl cell activity did not predict changes in delta power beyond 5 sec time window (Fig 2g; Supp Fig 7a). Cross-correlation analysis further revealed that coupling between calcium activity and delta power peaked at a short positive lag, consistent with near-simultaneous co-variation rather than predictive lead–lag relationships (Fig 2h). In addition, neither the mean activity of Sst-Chodl cells during SWS bouts nor other examined features were correlated with SWS bout duration (Supp Fig 7b-c). Together, these results suggest that Sst-Chodl cell activity reflects ongoing sleep-related cortical dynamics rather than anticipating forthcoming state transitions.

We examined how Sst-Chodl cell activity relates to sleep dynamics on a shorter timescale. Slow-wave sleep is not a homogeneous state but rather consists of rhythmic alternation between “DOWN” phases, characterized by near-complete suppression of spiking during upward deflections of the cortical LFP (DOWN states), and “UP” phases of elevated spiking activity (UP states; Fig 2i)^65^. Aligning Sst-Chodl cell activity to DOWN-state transitions revealed that these cells increased their activity preceding DOWN-state onset, whereas other neurons, simultaneously recorded as multi-unit spiking activity from the same recording site, exhibited heterogeneous responses with prominent rebound firing following DOWN-to-UP transitions (UP onset; Fig 2j-k and Supp Fig 7d). Notably, Sst-Chodl cell activity was not correlated with either the duration or amplitude of the subsequent DOWN state (Supp Fig 7e-f), suggesting that these neurons are preferentially engaged near the termination of cortical UP states rather than during quiescent DOWN phase itself.

To evaluate the specificity of Sst-Chodl cell activity patterns, we compared their state modulation to that of the broader Sst population and other neuronal classes using two publicly available datasets. First, using the Allen Brain Observatory dataset ^66^, we examined the correlations between Sst cell activity and arousal metrics, including locomotion and pupil area state (Supp Fig 8a). In this dataset, the Sst population as a whole was predominantly correlated with increased arousal (Supp Fig 8b-c). Next, using an independent dataset ^24^, we compared the activity of Sst-Chodl cells to other simultaneously recorded neuronal types (Supp Fig 8d-f). In contrast to arousal-activated interneuron subtypes, including regular-spiking Sst cells, Sst-Chodl cells showed negative correlations with arousal (Supp Fig 8d-f). Among IN populations, Sst-Chodl cells exhibited their strongest positive correlation with fast spiking Sst subtypes, particularly a numerically minor, Tac1 -expressing population (Supp Fig 8e-f).

Together, our results suggest that, unlike other Sst cell populations, Sst-Chodl cell activity closely mirror changes in behavioral/arousal states, becoming most active during highly synchronized cortical states.

### Activation of Sst-Chodl cells induces neocortical synchrony

To test whether Sst-Chodl cell activity causally promotes neocortical synchrony, we optogenetically activated these cells while performing high density electrophysiological recordings in head-fixed animals. We acutely inserted 64-channel linear silicon probes equipped with tapered optical fibers into the V1 of mice virally expressing ChR2 in Sst-Chodl cells (Fig 3a). This approach enabled simultaneous optogenetic stimulation of neocortical Sst-Chodl cells and recording of LFPs and single-unit activity across cortical layers (Fig 3b). Stimulation as delivered in ∼30 min blocks interleaved with non-stimulation (spontaneous) blocks (see Methods). Stimulation blocks were further subdivided into alternating stimulation and inter-trial intervals (ITI) epochs of 30 sec each (Fig 3c-d). We quantified multiple complementary metrics of cortical synchrony derived from LFP and spiking activity to compare network states with and without Sst-Chodl cell activation (Fig 3d-n).

**Figure 3.**
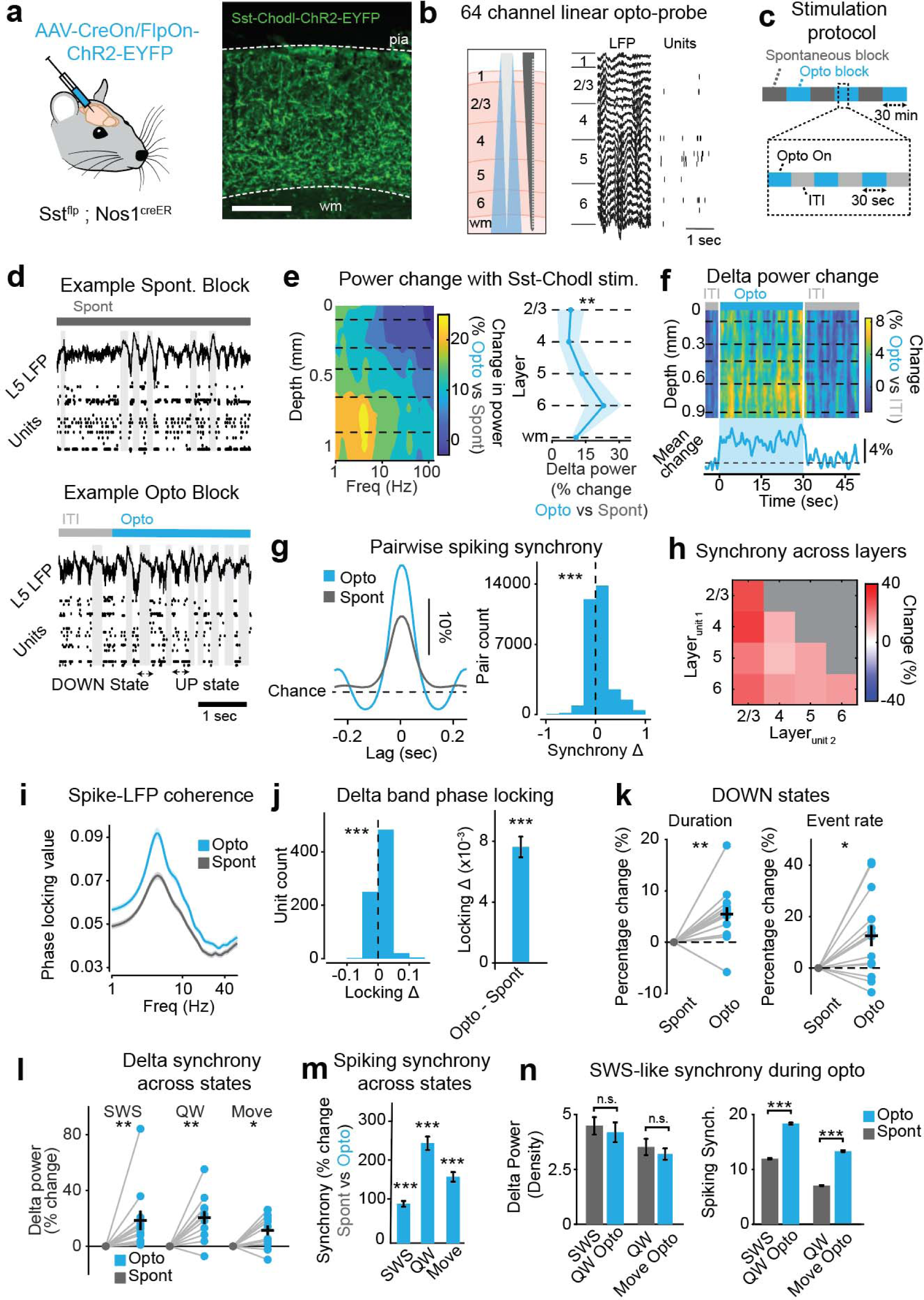
Optogenetic activation of Sst-Chodl cells induces robust neocortical network synchronization. **a)** Left: schematic of CreOn/FlpOn-ChR2-EYFP injections used to selectively express ChR2 in Sst-Chodl cells in V1. Right: example ChR2 expression (scale: 200µm). This approach enables activation of Sst-Chodl cells during electrophysiological recordings. **b)** Schematic of the 64-channel linear silicon probe with an attached tapered optical fiber, and example LFPs and isolated unitsaligned to cortical layer locations. **c)** Stimulation protocol consisting of spontaneous (Spont) and optogenetic stimulation (Opto) blocks. Opto blocks include repeated cycles of stimulation and inter-trial intervals (ITIs). **d)** Example trials showing LFPs from layer 5 and multi-unit activity during stimulation and ITIs; DOWN states are shaded. These traces illustrate that stimulation increases the occurrence and depth of synchronized events. **e)** Left: mean LFP power change across depths (Opto vs. Spont). Right: change after stimulation in delta band (1-4Hz) power during SWS across layers (p = 0.008, paired t-test). This demonstrates that Sst-Chodl cell activation enhances low-frequency throughout the cortical column. **f)** Average delta band power change across layers, with mean change below (mean Opto vs. ITI, p = 0.004, paired t-test). Stimulation rapidly increases delta power relative to the onset of the blue light. **g)** Normalized spike-train cross-correlograms (CCGs) during spontaneous and stimulated periods. Left: mean CCG. Right: distribution of CCG peak changes for all neuron pairs (p < 0.001, paired t-test). Stimulation increases spiking synchrony across the population. **h)** Change in pairwise spiking synchrony across cortical layers, showing that synchrony increases broadly and is not layer-specific. **i)** Spike-phase coherence across the LFP frequency spectrum during spontaneous and stimulation blocks. Sst-Chodl cell activation enhances coherence in the delta band. **j)** Distribution (left) and mean (right) of unit-wise change in delta-band phase locking with stimulation (p < 0.001, paired t-test). Spikes become more tightly locked to delta oscillations during stimulation. **k)** Increased DOWN-state duration and event rate during stimulation relative to spontaneous periods (p = 0.002 and p = 0.02, paired t-test). These changes indicate longer and more frequent synchronized OFF periods. **l)** Delta power change across SWS, quiet wake (QW) and movement (Move) (p = 0.008, 0.001, 0.008, paired t-test). Stimulation increases delta power across all states, even during movement where delta power is typically low. **m)** Change in pairwise spiking synchrony across SWS, QW, and Move (p < 0.001 for all, paired t-test). Synchrony increases across all states with stimulation. **n)** Comparison of delta power (density) and spiking synchrony (% above chance level) during spontaneous and stimulated conditions for SWS and QW (repeated measures ANOVA p < 0.001, with post hoc Tukey’s HSD; Delta: SWS-QW Opto p = 0.351, QW-Move Opto p = 0.700, SWS-Move Opto p = 0.003, SWS-QW p = 0.003, QW Opto-Move Opto p=0.034. Spiking synchrony: all comparisons p<0.001). Stimulation during QW increases delta power to SWS-like levels and elevates spiking synchrony beyond levels observed in spontaneous SWS. This shows that Sst-Chodl cell activation can drive synchrony to physiologically high, and even supraphysiological, levels. These experiments were performed in V1 of head-fixed mice. N = 14 sessions across 10 mice, n = 758 units and 32,710 pairs. ***: p < 0.001, **: p < 0.01, *: p < 0.05; ns, not significant. All Opto vs. spontaneous comparisons were made during SWS. Data are means; shading and bars indicate s.e.m.

**Figure 4.**
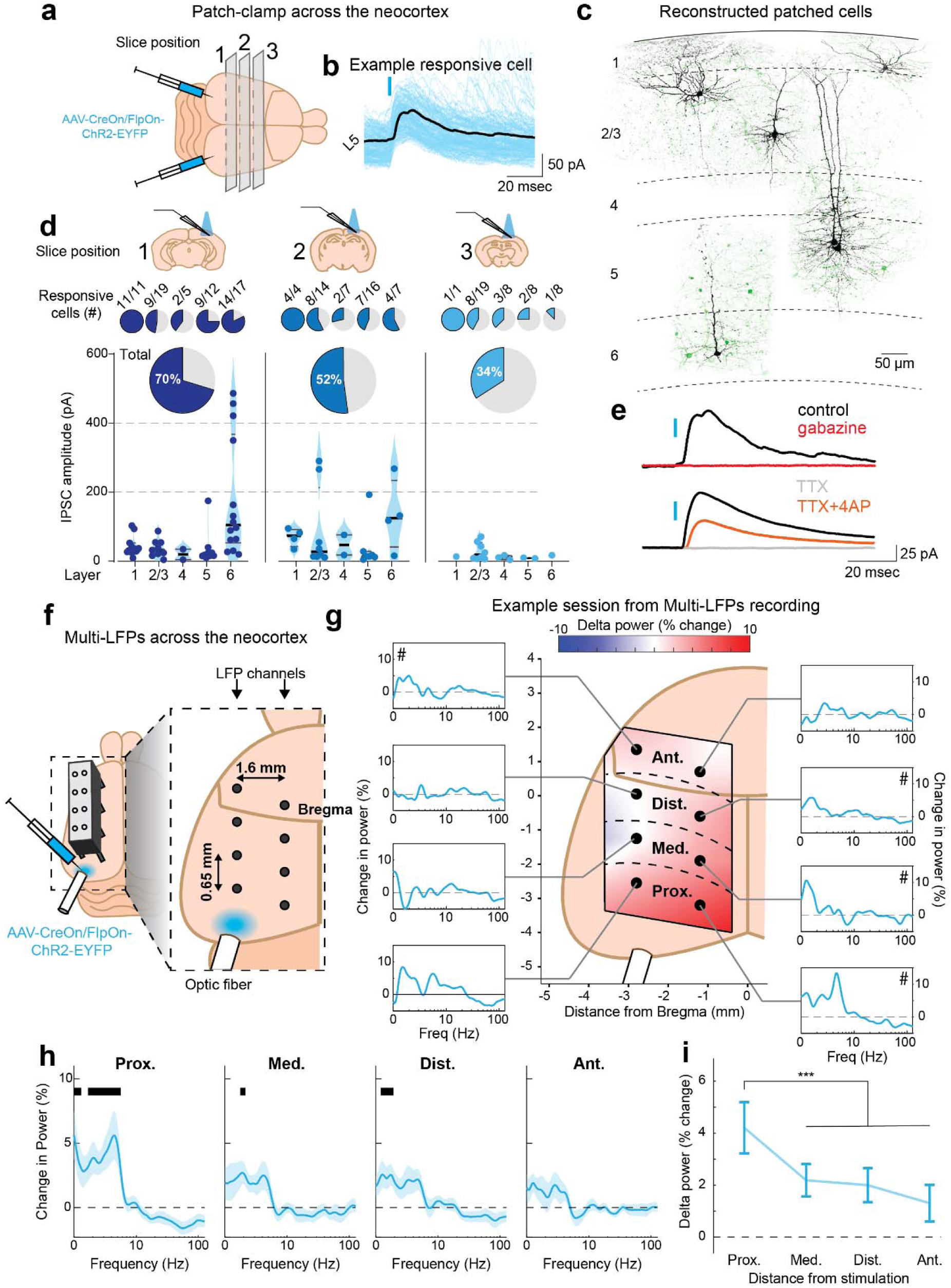
Sst-Chodl cells exert long-range inhibition and promote network synchronization across the neocortex. **a)** Illustration of the patch clamp approach for identifying long-range postsynaptic targets of Sst-Chodl cells following ChR2 expression in V1. **b)** Example optogenetically evoked inhibitory postsynaptic currents (oIPSCs; blue) and averaged response (151 traces; black) from a layer 5 pyramidal neuron aligned to the light stimulus (blue tick). These recordings confirm functional inhibitory synapses from Sst-Chodl axons. **c)** Schematic of cortical layers 1-6 near the injection site (V1) with representative anatomical fills of postsynaptic neurons (black) and ChR2-expressing Sst-Chodl fibers (green), highlighting the diversity of postsynaptic cell types encountered. **d)** Summary of responsive cells (blue) and oIPSC amplitudes across layer, grouped by anteroposterior distance from the injection site (antero-posterior distance from Bregma, |1|: -0.8 to -1.5; |2|: -1.5 to -2.2; |3|: -2.2 to -3.2; injection site is: -3.2mm). Horizontal lines show medians and interquartile range. Pie charts indicate the percentage of neurons responding to Sst-Chodl cell stimulation. Although responsiveness declines with distance, nearly one-third of cells recorded ∼2 mm from the injection site still showed oIPSCs, with layer 1 exhibiting the highest proportion of responsive neurons. Layer 6 responses showed the largest amplitudes. N = 16 animals (Sst^flp^ ; Nos1^creER^ mice), n = 156 cells (44, 48, and 64 cells per distance bin). **e)** Top: representative oIPSC from a layer 5 cell under control conditions (black) and after gabazine application (red; 50µM), confirming GABA_A receptor mediation. Bottom: TTX (0.5 µM), abolishing the oIPSC (grey), and addition of 4-AP (100 µM) restores it (orange), demonstrating that these responses are monosynaptic. **f)** Illustration of the multi-site LFP recording strategy used to measure long-range network effects, with electrodes positioned from ∼2 to 6 mm away from the V1 stimulation site. These experiments were performed on head-fixed mice. **g)** Example session showing stimulation-evoked changes in power spectra at each recording position (Proximal: ∼2.1mm; Medial: ∼3.2-3.5mm; Distal: ∼4.4-4.8mm; Anterior: ∼5.6-6.1mm from the injection site). # denotes the channel exhibiting the largest delta-band increase at each distance. **h)** Average power spectra from channels with the greatest delta increases at each distance. Black lines indicate frequency ranges with significant stimulation-induced changes (linear mixed-effects model with Bonferroni post hoc tests, p < 0.05). **i)** Average change in delta power (1-4Hz) as a function of distance from the stimulation site (repeated-measures ANOVA on linear mixed-effects model: distance effect p = 0.005; Proximal differs from Medial/Distal/Anterior, all p < 0.001; Medial vs. Distal vs. Anterior, p = 1). Sst-Chodl cell activation induces widespread low-frequency that diminishes with increasing distance from V1. N= 14 sessions from 7 mice. Data are means; shading and bars indicate s.e.m.

We first, examined changes in LFP power across cortical depth during stimulation relative to spontaneous periods. Optogenetic activation of Sst-Chodl cells led to a robust increased in delta-band power, hallmark of synchronized cortical states, across all layers, with the largest effects observed in deeper layers (Fig 3e). This increase emerged rapidly following stimulation onset (Fig 3f). We next assessed spiking synchrony among well-isolated single units (putative single neurons) by computing spike train cross-correlations. Sst-Chodl cells activation increased the magnitude of spiking synchrony (Fig 3g, Supp Fig 9a), an effect observed across all layers (Fig 3h), and independent of the stimulation frequencies (Supp Fig 9b) or unit class (regular spiking versus fast spiking) (Supp Fig 9c). Although Sst-Chodl cell activation produced a slight reduction in overall firing rate, spike train structure and burst properties were largely preserved (Supp Fig 9d-h), indicating that enhanced synchrony did not result from increased firing but instead reflected changes in temporal coordination across neurons.

To determine whether Sst-Chodl cell-evoked synchrony extended spatially within the visual cortex, we used multishank 64-channel silicon probes comprising six shanks spaced 200 µm apart, with the most lateral shank coupled to a tapered optical fiber (Supp Fig 10a). Single-shank stimulation reliably increased delta power and spiking synchrony at the stimulated site and, interestingly, these effects propagated across all shanks, spanning ∼1 mm of cortex (Supp Fig 10b-c). This spatial spread indicates that Sst-Chodl cell activation induces distributed synchronization across visual cortical networks.

Given the close relationship between LFP fluctuations and spiking activity, we next assessed spike–field coupling during stimulation by quantifying spike-phase locking values. Under baseline conditions, spikes were preferentially locked to the delta phase of the LFP (Fig 3i), and Sst-Chodl cell stimulation strongly enhanced this coupling (Fig 3i-j, Supp Fig 9i). As an extreme manifestation of phase locking, DOWN states, brief periods of spiking silence which are prominent during SWS, became more frequent and longer-lasting during stimulation, with both event rate and duration significantly increased (Fig 3k, Supp Fig 11a-d).

We then asked whether Sst-Chodl cell activation modulates cortical synchrony across different states. Comparing stimulation epochs to matched spontaneous periods revealed significant increases in delta-band power during SWS, Quiet Wake, and Movement (Move; Fig 3l, Supp Fig 11e-i; REM was excluded due to insufficient bout duration). Spiking synchrony was similarly enhanced across all three states (Fig 3m, Supp Fig 9j-k). Notably, synchrony increased even during Movement, when Sst-Chodl cells are typically not active endogenously (Fig 3l-n).

Finally, we compared the magnitude of stimulation-induced synchrony to physiological levels observed during spontaneous SWS and QW (Fig 3n, Supp Fig 9j-k, Supp Fig 11g-h). Sst-Chodl cell activation during QW elevated delta power to levels comparable to spontaneous SWS, whereas stimulation during movement produced delta power comparable to that observed in spontaneous QW (Fig 3n; Supp Fig 11g–h). Notably, Sst-Chodl cell activation in either state produced spiking synchrony that exceeded the maximum levels observed during spontaneous SWS (Fig 3n, Supp Fig 9j-k), underscoring the strong synchronizing capacity of this cell population.

Taken together, these findings demonstrate that activation of Sst-Chodl cells robustly enhances neocortical synchrony, recapitulating, and in some cases exceeding, key features of endogenous synchronized cortical states observed during SWS.

### Stimulation of non-Chodl Sst cells does not increase cortical synchrony

Next, we examined whether activation of a non-Chodl subtype of Sst interneurons is sufficient to induce cortical synchrony. To address this, we selectively expressed ChR2 in Martinotti cells,^67^ a major population of non-Chodl Sst cells, using an enhancer-driven viral strategy ^67^ and recorded cortical activity during optogenetic stimulation (Supp Fig 12a). As expected, Martinotti cell stimulation produced a reduction in overall cortical firing rate (Supp Fig 12b). However, in contrast to Sst-Chodl cell activation, this manipulation did not alter cortical synchrony, as measured by delta-band LFP power or spiking synchrony (Supp Fig 12c-d). These findings indicate that the capacity to promote neocortical synchrony is not shared across all Sst INs but is instead a distinguishing feature of the Sst-Chodl cells.

### Sst-Chodl cells regulate the activity and synchrony in distant neocortical areas

We have shown that Sst-Chodl cells are long-range projecting neurons that broadly target neocortical areas (Fig 1) and that their activation promotes regional network synchronization within the visual cortex (Fig 3, Supp Fig 10). We next asked whether Sst-Chodl cells can also influence neuronal activity and cortical synchrony at more distant neocortical sites.

To directly determine the spatial extent of their synaptic targets, we expressed ChR2 in Sst-Chodl cells in V1 and performed whole-cell patch-clamp recordings from putative postsynaptic neurons in acute neocortical slices, at locations ranging from the injection site to ∼2.4 mm away (Fig 4a-b, Supp Fig 13). Recordings were grouped by distance from the injection site (see Methods) and optogenetically-evoked inhibitory postsynaptic currents (oIPSCs) were assessed in response to brief blue-light pulses (0.5 msec, 473 nm). Postsynaptic targets of Sst-Chodl cells included both pyramidal neurons and interneurons (Fig 4c, Supp Fig 13b). Within the V1 (injection site), oIPSCs were observed in the majority of recorded neurons, with particularly high response rates in layers 1, 5 and 6 (Fig 4d, Supp Fig 13b-h). Although the proportion of responsive cells decreased with increasing distance (Chi squared including Layer 1 cells p < 0.001; Chi squared excluding Layer 1 cells p = 0.009), oIPSCs remained detectable in nearly one-third of neurons recorded approximately 2 mm from the injection site (Fig 4d). Consistent with anatomical tracing showing prominent long-range axons in Layer 1 (Fig 1h, Supp Fig 3c), this layer exhibited the highest proportion of responsive cells across all recording sites (Fig 4d, Supp Fig 13b). Response amplitudes also varied by layer, with significantly larger oIPSCs observed in Layer 6 compared to other layers (Fig 4d, Supp Fig 13b-d). Finally, we confirmed that these oIPSCs were GABAergic and monosynaptic: responses latencies were short, responses persisted in presence of tetrodotoxin and 4-aminopyridine (0.5 and 100 µM), and currents were abolished by the GABA _A_ receptor antagonist gabazine (50 µM; Fig 4e, Supp Fig 13e-f). Together, these results align with our anatomical data and demonstrate that Sst-Chodl cells provide direct, long-range inhibitory across neocortical layers.

Given the broad ipsilateral intracortical projections of Sst-Chodl cells, we next asked how far the synchronization induced by their activation propagates across the neocortex. To address this, we expressed ChR2 in Sst-Chodl cells in V1 and recorded LFPs from eight ipsilateral neocortical locations positioned well beyond the spatial extent sampled in our multishank recordings (Supp Fig 10), spanning distances of approximately 2-6 mm from the stimulation site (Fig 4f). Using a similar optogenetic stimulation protocol, we compared LFP power spectra during stimulation and ITI periods at each recording location (Fig 4g), and quantified changes in power as a function of distance from the stimulation site (see Methods). Optogenetic activation of Sst-Chodl cells in V1 increased low-frequency LFP power across most of ipsilateral recording sites, extending from visual cortex to frontal motor areas (Fig 4g-i). This effect was strongest at sites closest to the stimulation region and progressively declined with increasing distance (Fig 4i). These results demonstrate that Sst-Chodl cell activation can drive neocortical synchronization over millimeter-scale distances within the ipsilateral cortex, with a graded decay consistent with long-range intracortical influence.

### Neocortical activation of Sst-Chodl cells promotes sleep

Previous work has suggested that during periods of intense SWS, characterized by prominent cortical synchrony, putative Sst-Chodl cells are active throughout the neocortex ^46^. Because stimulation of Sst-Chodl cells induced a synchronized neocortical state resembling SWS (Fig 3-4 and Supp Fig 10), we hypothesized that widespread neocortical activation of these cells might not only enhance cortical synchrony but also promote behavioral features of sleep. To test this hypothesis, we utilized an INTRSECT-based chemogenetic approach and bilaterally injected an AAV8-CreOn/FlpOn-hM3Dq-mCherry vector into 22 neocortical locations (11 injections per hemisphere) to selectively express an excitatory DREADD in neocortical Sst-Chodl cells (Fig 5a, Supp Fig 14). Using 2-photon imaging, we confirmed that systemic administration of a low dose of the DREADD agonist, clozapine-N-oxide (CNO; 0.5 mg/kg), increased Sst-Chodl cell activity largely independently of the mice’s arousal state (Supp Fig 14b-c, Supp Movie 2-3). To test the impact of pan-neocortical Sst-Chodl cell activation on sleep behavior, we performed counterbalanced systemic injections of vehicle or CNO and monitored LFP activity and movement in freely behaving mice in their home cage for 2 hours during the light (inactive) phase of their circadian cycle (Supp Fig 14d). Chemogenetic activation of Sst-Chodl cells increased the duration of both SWS and REM sleep relative to vehicle treatment (Fig 5b-e, Supp Fig 14e). This manipulation also reduced sleep latency and increased the number of SWS bouts, while leaving bout duration unchanged (Fig 5f-g, Supp Fig 14f).

**Figure 5.**
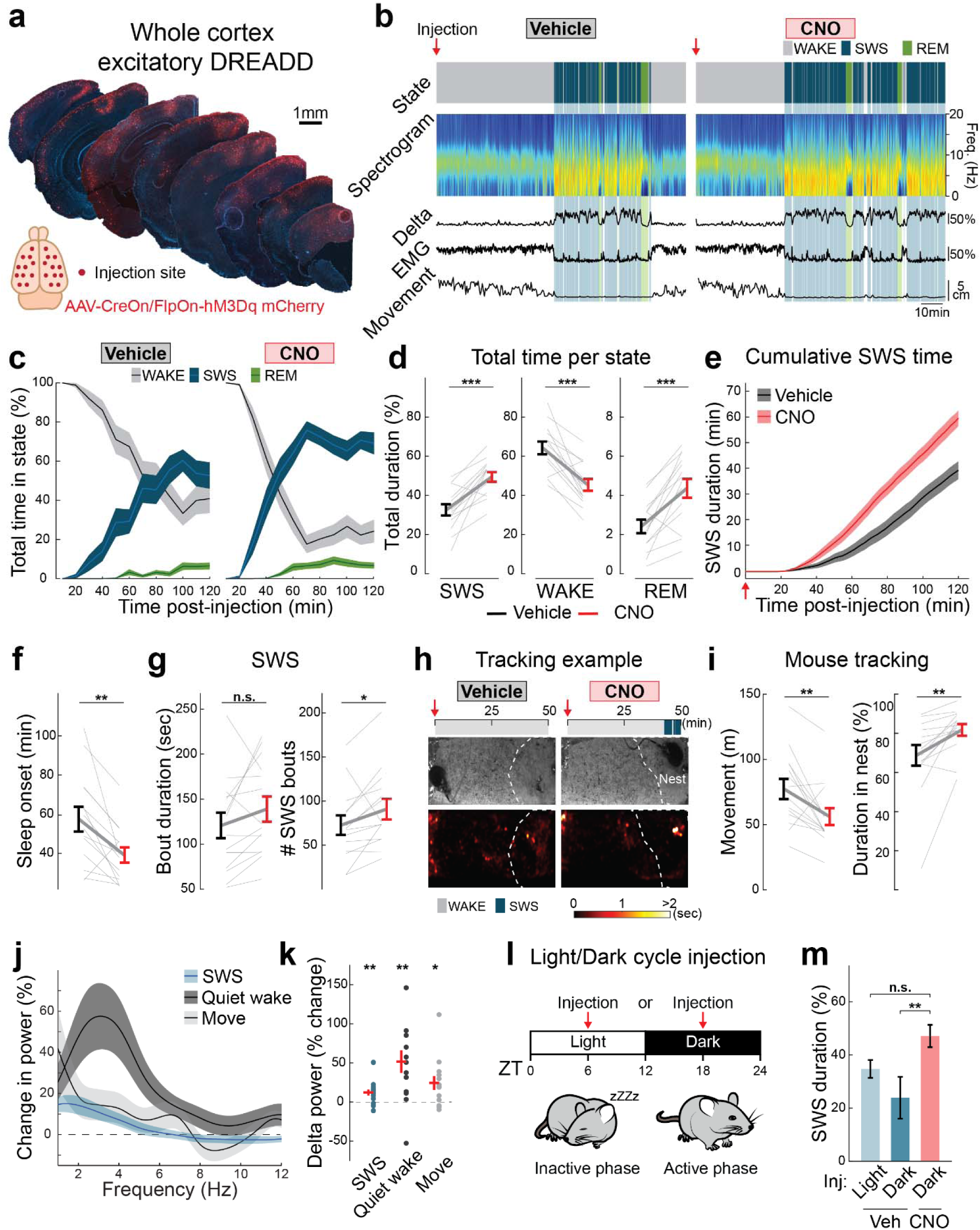
Chemogenetic activation of neocortical Sst-Chodl cells promotes sleep. **a)** Strategy for global activation of Sst-Chodl cells: an INTRSECT AAV-EF1a-fDIO-mCherry vector was injected at 22 neocortical sites (11 per hemisphere). Right: control brain showing broad expression of the CreOn/FlpOn-mCherry reporter. This approach enabled pan-neocortical recruitment of Sst-Chodl neurons. **b)** Example of freely moving recordings during the light/inactive phase for one mouse following systemic injection of Vehicle (left) or CNO (0.5 mg/kg; right), showing sleep/wake states and associated physiological measures. CNO increased time spent in sleep states. **c)** Percentage of time spent in WAKE, SWS and REM after Vehicle or CNO injection. **d)** Total time in each state across the 2-hours post-injection window (SWS, p < 0.001; WAKE, p < 0.001; REM, p < 0.001, paired t-test). CNO increased both SWS and REM while reducing Wake. **e)** Cumulative SWS time across sessions. **f)** Latency to sleep onset (p = 0.003, paired t-test). **g)** SWS bout duration (ns, p = 0.110, paired t-test) and numbers of SWS bouts (p = 0.036, paired t-test). These data show that Sst-Chodl cell activation promotes sleep primarily by increasing bout number and reducing latency, rather than lengthening individual bouts. **h)** Example animal tracking from the first 50 minutes after injections. Top: sleep/wake scoring. Middle: mouse position and nest location (white dashed line). Bottom: heatmap of spatial occupancy. CNO-treated animals spent more time resting in the nest. **i)** Total distance travelled (p = 0.005, paired t-test) and time in nest (p = 0.003, paired t-test) over 2 hours post-injection. Reduced locomotion and increased nest occupancy mirror the behavioral shift toward sleep. **j)** LFP power changes recorded in V1 during SWS, quiet wake (QW), and movement (Move). **k)** Delta-band (1-4Hz) power changes across states (SWS p = 0.008; QW p = 0.004; Move p = 0.018; paired t-tests). Chemogenetic activation increases low-frequency across arousal states, consistent with the effects of optogenetic stimulation. **l)** Illustration of the experimental timing when treated animals during light/inactive (ZT3–7) vs dark/active (ZT18) phase. **m)** Comparison of SWS duration across light/inactive vs. dark/active conditions and Vehicle vs CNO (ANOVA on mixed-effects model, group effect p = 0.004; Bonferroni post hoc: Light Veh vs Dark Veh p = 0.214, Light Veh vs Dark CNO p = 0.100, Dark Veh vs Dark CNO p = 0.003). CNO increased SWS in both phases, driving sleep even during the normally wake-dominant dark/active period. These findings demonstrate that global activation of neocortical Sst-Chodl cells is sufficient to enhance cortical synchrony and strongly promote sleep across behavioral contexts and circadian phases. N = 14 mice (Light Veh = 16 sessions; Dark Veh = 7; Light CNO = 7). *** p < 0.001, ** p < 0.01, * p < 0.05, ns = not significant. Injections were performed during the light phase for b-m. Data are means; shading and bars indicate s.e.m.

To evaluate whether these changes were accompanied by typical sleep-associated behavior, we used DeepLabCut ^68^ to quantify locomotion and nest occupancy following global activation of neocortical Sst-Chodl cells. CNO-treated mice traveled shorter distances and spent more time in the nest (Fig 5h-i, Supp Fig 14g), consistent with a reduced proportion of time spent in movement relative to quiet wakefulness (Supp Fig 14h-i) and with the observed increase in sleep (Fig 5b-g).

We next examined the effects of pan-neocortical Sst-Chodl cell activation on LFP dynamics. Similarly to optogenetic manipulation (Fig 3), chemogenetic activation of the neocortical Sst-Chodl cells increased neocortical LFP power predominantly in low-frequency bands and largely independently of behavioral state (Fig 5j-k, Supp Fig 15a). This effect was prominent in neocortical recording sites but minimal in hippocampal channels (Supp Fig 14j-k, Supp Fig 15b), indicating a preferential impact on neocortical circuits. Importantly, CNO administration in control mice lacking DREADD expression did not alter sleep sleep-wake architecture (Supp Fig 15c-f) or LFP power (Supp Fig 15g).

Because the initial behavioral experiments were performed during the light (inactive) phase, when mice naturally spend more time asleep, we next tested whether Sst-Chodl cell activation would exert similar effects during the dark (active) phase. As expected, vehicle-treated mice during the dark (active) phase (Zeitgeber Time, ZT18) spent more time awake and exhibited longer sleep latencies than during the light (inactive) phase (Fig 5m, Supp Fig 16). Nevertheless, CNO administration during the dark (active) phase increased SWS duration and reduced sleep latency relative to vehicle treatment (Supp Fig 16b-g). Notably, sleep amounts induced by Sst-Chodl cell activation during the dark (active) phase were comparable to, and in some cases exceeded, baseline sleep observed during the light (inactive) phase of vehicle treated animals (Fig 5m), indicating a robust sleep-promoting effect.

Together, these results indicate that, independent of circadian phase, global activation of neocortical Sst-Chodl cells is sufficient to promote cortical synchrony and sleep.

## Discussion

Here, we show that long-range, neocortical Sst-Chodl cells – a unique neuronal subtype ^30,36^ that is highly conserved across evolution ^33,41,42^ – are selectively active during low arousal synchronized states and promote patterns of synchronous neocortical activity. We propose that Sst-Chodl cells synchronize neocortical networks through their extensive local and long-range projections, positioning them as a key cortical component in the regulation of sleep.

Cortical synchronization states are generated by the interactions of a variety of cell types and circuit components within and outside of the cortex. In particular, previous work has shown that cortical fast-spiking parvalbumin-expressing cells are critical for the generation of desynchronized states by enhancing gamma oscillations ^26,27^. In contrast, we show that the activity of Sst-Chodl cells not only reflects cortical synchronization states but also actively promotes them, enhancing delta-frequency LFP fluctuations and coordinated spiking across neuronal populations. Sst-Chodl cells exhibit exceptionally dense projections and high synaptic connectivity across all cortical layers within the cortical region where their cell bodies are located, making them well suited to promote local cortical synchrony. This feature may be particularly relevant for forms of local sleep, in which circumscribed cortical regions exhibit sleep-like patterns in the absence of overt behavioral sleep ^69^. In addition, we demonstrate that Sst-Chodl cells extend long-range axons that inhibit neurons millimeters away from their somata, suggesting a mechanism by which these cells could contribute to synchronization across distant neocortical regions ^70^.

The prevailing framework for sleep regulation emphasizes subcortical structures as primary drivers of sleep initiation and maintenance, with the cortex viewed largely as a passive downstream recipient ^71^. However, accumulating evidence supports a more active role for the neocortex in sleep-wake regulation. In particular, a subset of cortical Sst-expressing cells have been implicated in sleep promotion and sleep-preparatory behaviors ^54,55^. Our findings extend this framework by identifying Sst-Chodl cells as a distinct cortical population capable of shaping neocortical synchrony and sleep-related dynamics. Future studies will be required to determine whether Sst-Chodl cells outside sensory cortices receive inputs from, or project to, canonical sleep-promoting regions in the hypothalamus or midbrain, or whether their influence on sleep is mediated primarily through intracortical mechanisms. Notably, prior work has shown that cortex-wide reductions in synaptic output from deep-layer pyramidal neurons attenuate homeostatic sleep rebound ^56^ indicating that cortical output can contribute to sleep regulation. In this context, we find that Sst-Chodl neurons provide particularly strong synaptic inhibition and LFP modulation in deep cortical layers, where pyramidal neurons associated with synchronized states are concentrated ^72–74^. These observations raise the possibility that Sst-Chodl cells influence sleep homeostasis by shaping the activity of deep-layer cortical circuits. Supporting this notion, previous *ex vivo* studies have implicated putative Sst-Chodl cells in homeostatic recovery from sleep deprivation ^45,46^, a relationship that remains incompletely understood and warrants further investigation.

Although cortical Sst-expressing INs as a group have been implicated in the regulation of cortical states and linked to sleep-related processes ^54,55^, their functional roles have been difficult to reconcile across studies, with reported effects spanning gamma ^75,76^, beta ^77^, theta ^78^, and delta rhythms ^54^. Single-cell transcriptomic profiling has revealed substantial heterogeneity within the Sst population, indicating that distinct Sst subclasses may subserve divergent, and potentially opposing, functions ^24,32^. In this context, differences in experimental approaches may preferentially engage different Sst subtypes, contributing to variability in reported outcomes. Consistent with this interpretation, we identified a minor population of cells within the Sst-Nos1 intersection that were anti-corelated with arousal and likely correspond to non-Sst-Chodl, Type II Nos1 cells expressing low levels of Sst and Nos1 ^32,63^. These cells may participate in neurovascular control during movement ^37,43,79^, consistent with a functional role distinct from the promotion of synchronized cortical states. Our results highlight the importance of transcriptomically informed targeting strategies for dissecting IN subclass function and underscore the utility of this approach for understanding cortical circuit mechanisms.

Beyond their synaptic inhibitory effects, Sst-Chodl cell express additional signaling molecules that may contribute to their function. Notably, these neurons likely represent the largest source of nitric oxide (NO) signaling in the neocortex ^48^. Mice with reduced Nos1 exhibit disrupted sleep architecture, including reduced ability to sustain prolonged SWS bouts and impaired homeostatic response to sleep deprivation ^45^. Similarly, selective deletion of Nos1 from Sst cells results in deficits in delta rhythms ^80^. Potential targets of NO signaling from Sst-Chodl cells include parvalbumin-expressing INs ^48,81^ and components of the neurovascular system ^37,79^. The slow, synchronized electrical activity promoted by Sst-Chodl cell activation may therefore be mirrored in vascular dynamics, potentially supporting rhythmic fluid movement during glymphatic clearance in sleep ^82,83^.

The mechanisms governing the state-dependent activation and suppression of Sst-Chodl cells remain to be elucidated. While thalamocortical circuits are well known to regulate cortical UP/DOWN state dynamics ^17^, our monosynaptic tracing experiments indicate that Sst-Chodl cells receive predominantly neocortical inputs, with minimal thalamic contributions. Prior studies have shown that putative Sst-Chodl cells receive extensive neuromodulatory inputs^48,84,85^ which may not be captured by our rabies-based tracing approaches. Our finding that Sst-Chodl cell activity increase prior to DOWN state onset suggests that these neurons may participate in regulating UP/DOWN transitions, similar to other inhibitory neuron classes^86–88^. Activation of Sst-Chodl cells could arise through disinhibition mediated by Vip expressing INs, consistent with our rabies tracing data indicating Vip inputs onto these cells, or through changes in arousal-related neuromodulatory tone, such as acetylcholine ^84^. This interpretation is consistent with previous studies showing that Sst-Chodl cells strongly express inhibitory muscarinic M2 and M4 receptors, as well as inhibitory 5HT1a receptors ^30,48,84,89^. Another possibility is that sleep-promoting substances (somnogens), such as adenosine, could activate these cells during synchronized cortical states ^43^. In addition, Substance P has been shown to activate Sst-Chodl cells, and local application of Substance P promotes cortical slow-wave activity and electrophysiological signatures of SWS ^44,80,84^, although its precise role in sleep regulation remains to be elucidated ^85^. In this context, we find that Sst-Chodl cells receive input from both Pvalb- and Sst-expressing INs, which represent potential sources of Substance P. Identifying additional G-protein coupled receptors expressed by Sst-Chodl neurons will be important for elucidating the signaling pathways through which sleep- and arousal-related factors engage this cell population.

Overall, this work identifies Sst-Chodl cells as a key cortical circuit element associated with low-arousal neocortical states and sleep. The properties of this genetically distinct IN subtype point to an active contribution of cortical circuits to sleep-related dynamics, complementing established roles of subcortical structures in sleep regulation. Together with prior studies, our findings highlight an underappreciated role for the cortex in shaping sleep states. Given that sleep disturbances are a prominent feature of many neuropsychiatric, neurodevelopmental and neurological disorders, this highly conserved cell population may represent a useful entry point for future mechanistic and translational investigations.

## Acknowledgements.

We would like to thank Eliezyer Fermino de Oliveira for help using buzcode tools, Kevin Fisher and Kostantin Dobrenis from the Einstein Neural Cell Engineering and Imaging Core and Hillary Guzik and Vera Desmarais for assistance with imaging and image analysis, Andrea Velez, Cecilia Cuddy, Alina Widmann, Hikari Tanaka and Neel Patel for providing morphological reconstruction annotations and tissue cutting, Gabriel Batazar for helping in building LFP implants, Josh Siegle and Anna Lakunina for performing experiments not present in the manuscript, Alexandra Quezada for assistance with flexible probe methods, Julia Andrade, Ingrid Redford, Rachel Dalley, Sarah Walling-Bell, Clare Gamlin and Olga Gliko for helping in single cell reconstruction and Adam Kohn, Saleem Nicola, Tiago Gonçalves, and Jean Hebert for helpful guidance during these studies.

This work was supported by a NARSAD Young Investigator Award, a Simons Bridge to Independence Award, a Whitehall Award, and National Institutes of Health (NIH) grants R01EY034617, R21MH133097, R01EY034310 to RBB, by NINDS F31NS120723 to JMR, by Leon Levy Scholarships in Neuroscience 2023 20203010, NARSAD Young Investigator Award 31835 and Philippe Foundation to GT, by NIH 5R01HL059658 to TSK, by NIH 1U19MH114830-01 to H.Z, by Clayton Foundation for Research, W.M. Keck Foundation, BD2 Foundation, WoodNext Foundation, the NSF (24323797), and the Waggoner Center for Alcohol and Addiction Research to LEF, by P30CA013330/SIG31S10OD034397-01 to Hillary Guzik and Vera Desmarais and by NIH 1S10OD030508 to Kostantin Dobrenis, Einstein NCEI core.

## Author contributions

Conceptualization: JMR, GT, RBB. In vivo imaging: JMR and GT. Single cell reconstructions and analysis: AM, MM, BS, SS, JMR. Bulk morphological reconstructions: JMR, JM, BS. Ex vivo patch clamp: AVL, SL, SR. Headfixed optogenetics of Sst-Chodl cells: JMR, GT. Headfixed optogenetics of Martinotti cells: AVL, SK, JMR, LS. Freely moving chemogenetics: GT, JMR. Rabies staining/quantification: NPC, GM. Surgery and brain processing: JMR, GT, AVL, GNS, JM. Reagents: CR, LEF, BT, TD, DS, HZ, KD, JN. Scientific Advising: TK. Supervision: RBB. Initial draft: JMR, GT, RBB. All authors edited and approved the manuscript.

## Materials & Correspondence

Correspondence and requests for materials should be addressed to Renata Batista-Brito, (R.B.B) and Geoffrey Terral, (G.T).

## Materials and Methods

### Mice

All animal handling and maintenance was performed according to the regulations of the Institutional Animal Care and Use Committee of Albert Einstein College of Medicine (Protocol # 00001393). Sst^flp^ ^+/+^ ; Nos1^creER^ ^+/-^ and Sst^flp^ ^+/-^ ; Nos1^creER^ ^+/-^ animals were used interchangeably in this study with no detected differences between the two groups (Nos1^creER^: Jax # 014541, Sst^flp^: Jax # 031629). Nos1^creER^ ^+/-^ animals were crossed with Sst^flp^ ^+/+^ animals to obtain Sst^flp^ ^+/-^ ; Nos1^creER^ ^+/-^ animals and Sst^flp^ ^+/-^ ; Nos1^creER^ ^+/+^ were crossed with Sst^flp^ ^+/+^ animals to obtain Sst^flp +/+^ ; Nos1^creER^ ^+/-^ animals. Sst^flp^ ^+/+^ ; Nos1^creER^ ^+/-^ animals were crossed with Ai210 ^+/+^ animals (gift from Allen Institute)^90^, to obtain animals for *in vivo* imaging experiments. Chodl^cre^ ^+/+^ animals (gift from Allen Institute) ^90^ were crossed with Sst^flp^ ^+/+^ or Sst^flp^ ^+/-^ animals. Both adult male and female mice above P50 were used in this study. Animals were kept under a 12h light/dark cycle (lights on 7am or lights on 4pm for ZT18 chemogenetic experiments) under standard housing conditions.

### Cell type classification and sequencing

Data retrieved from ^90^.

### Surgical procedures

Mice were anesthetized with isoflurane (5% by volume for induction and between 1 and 2% for maintenance), placed on a stereotaxic frame, and kept warm with closed loop heating pad. For pain management, animals were given intraperitoneal (i.p.) meloxicam at 2.5 mg/kg and local lidocaine on the scalp before skin incision. The following coordinates are distance from Bregma and DV values refer to brain surface.

For viral injections, we minimized brain damage by performing burr holes, keeping a thin layer of the bone intact where the glass micropipettes could penetrate.

For wide labeling morphological reconstructions, a viral vector injection of 50 nL was injected into a burr hole split across two levels (DV -0.25 and DV -0.55) at AP -3.3, ML ±2.7 to target primary visual cortex (V1) or at AP -1, ML ±3 to target primary somatosensory cortex (S1).

For silicon probe coupled optogenetic experiments, four burr holes were drilled centered over V1 (AP -3.3, ML ±2.7) spaced ∼1mm apart, and positioned to avoid blood vessels visible through the skull. 500nL of virus was injected into each burr hole split across two levels (DV - 0.25 and DV -0.55) for a total of 2µL of virus.

For calcium imaging related surgeries, animals were also administered with dexamethasone at 4mg/kg 1h prior craniotomy. A 3mm craniotomy was made over V1, and if applicable, virus was injected into the center of this craniotomy while the brain was kept moist with hydrated gelatin surgical foam. Then a cranial window was placed over the opening and sealed to the skull with cyanoacrylate glue. For imaging Sst-Chodl cells in superficial layers, the cranial window consisted of a stack of 3 coverslips: two 3mm #1 glass coverslip (Warner Instruments) attached with Norland Optical Adhesive #71 to a 5mm glass coverslip allowing the 3mm glass to sit against the dura discouraging bone regrowth ^91^. For imaging Sst-Chodl cells in deep layers, durotomy was performed before implantation of a 1.5mm microprism (A1 coated, 4531-0023, Tower Optical Corporation) attached with Norland Optical Adhesive to a single 3mm glass coverslip.

For pan-neocortex viral delivery, 22 burr holes were split bilaterally and 400nL of viral vectors were injected at the following coordinates: AP +1,ML ±3, DV -1; AP +1, ±1.8, DV -0.4; AP -0.5, ML ±3.8, DV -1.5; AP -0.5, ML ±2.6, DV -0.4; AP -2, ML ±4, DV-1.2; AP -2, ML ±3.2, DV -0.3; AP -2, ML ±1.2, DV -0.3; AP -3.5, ML ±3.5, DV -0.3; AP -3.5, ML ±2, DV -0.3 and 800nL of viral vectors were injected at the following coordinates: AP 2.5, ML ±1.2, DV -1; AP 0, ML ±1, DV - 0.5.

For rabies tracing experiments, a total volume of 500nL of the helper mixture (2:1 ratio oG over TVA) split across two levels (DV -0.25 and DV -0.55) was injected into a single burr hole at AP - 3.3, ML ±2.7. Following a minimum of four weeks to allow robust expression of the helper constructs, rabies vector was delivered as a single injection of 150nL at the same AP/ML coordinates and DV -0.4. Mice were perfused 10 days after rabies delivery to capture the full extent of monosynaptic retrograde labeling while minimizing cytotoxicity.

For calcium imaging and whole-neocortex chemogenetic experiments, a separate burr hole was made in contralateral V1 for insertion of a made-in-house microwire array connected to a 16-channel omnetic (A79038-001, Omnetics connector corporation), used for measuring local field potentials (LFP), was inserted. The LFP wires consisted of 5 to 8 tungsten wires of 50µm spanning from middle layers of V1 area into CA1. For calcium imaging combined with flexible 32-channel probes (tetrode configuration, Neuralthread, ^92^), the electrode attached to a stainless-steel pole with adhesive polyethylene glycol was inserted perpendicular to the brain at the border of the craniotomy. Then, the polyethylene glycol was dissolved by applying saline continuously for 10min on top of the pole to ensure its disengagement from the electrode. After pulling the pole straight up out of the brain, the cranial window was positioned above the electrode.

For multisite LFP and optogenetic experiments, eight burr holes were made at the following coordinates: Anterior: AP +1.35, ML -2.8 and AP +0.7, ML -1.2; Distal: AP +0.05, ML -2.8 and AP -0.6, ML -1.2; Medial: AP -1.25, ML -2.8 and AP -1.9, ML -1.2; Proximal: -2.55, ML -2.8 and -3.2, ML -1.2. A made-in-house microwire array consisting of eight tungsten wires of 200µm aligned to the mouse skull was connected to a 16-channel omnetic and inserted in the surface of the brain. 500nL of virus was injected ipsilateral of the multi-LFPs at AP -2.7, ML -2.7 split across two levels (DV -0.25 and DV -0.55) and a fiber optic cannulae of 400µm diameter and 6mm long was positioned at the entrance of the burr hole.

Injections were performed using a Nanoject III system at a rate of 1-2nL/secs through glass micropipettes that were pulled and then ground to bevel with 40µm diameter using a Naragishe diamond wheel. For LFPs and flexible probes, a 200µm tungsten ground wire attached to a gold pin was inserted into the cerebellum, and a headpost was implanted as previously described ^93^.

All implants were secured to the skull using Optibond or Super-Bond followed by dental cement. When Nos1^creER^ ^+/-^ animals were used, tamoxifen (Thermo Scientific) was administered at least two weeks after surgery. A stock solution of 20mg/mL tamoxifen in corn oil was injected i.p. at 0.1mg tamoxifen/g body weight for five consecutive days. Mice were allowed for a minimum of four weeks for CreER-mediated recombination before experimental use.

The following AAV viral vectors were used in this study at concentrations between 10e12 - 10e13 vg/ml:

| **AAV vector** | **Serotype** | **Source** |
| --- | --- | --- |
| Ef1a-CreOn/FlpOn-oScarlet | 8 | provided by the lab of K. Deisseroth |
| Ef1a-CreOn/FlpOn-GCaMP6m | 8 | provided by the lab of K. Deisseroth |
| hSyn-CreOn/FlpOn-ChR2-EYFP | DJ | UNC vector core |
| EF1a-fDIO-mCherry | 5 | addgene (#114471) |
| nEF-CreOn/FlpOn-hM3Dq-mCherry | 8 | provided by the lab of K. Deisseroth |
| nEF-CreOn/FlpOn-TVA-mCherry WPRE | 8 | GVVC (#AAV-197) |
| Ef1a-CreOn/FlpOn-oG WPRE | 8 | GVVC (#AAV-198) |
| EnvA-DG-Rabies-EGFP | N/A | Salk vector core |
| iSSTe4-ChR2-mCherry<sup>67</sup> | 1 | provided by the lab of G. Fishell |

### Immunohistochemistry

Animals were perfused transcardially with ice-cold 4% paraformaldehyde (PFA) and brains were dissected and post-fixed in 4% PFA for 1 hour at 4°C. After sucrose cryoprotection, brains were embedded in OCT, cryosectioned at 20µm, and adhered to glass slides. For staining, sections with incubated in a blocking solution containing 1.5% Donkey serum, 1% Triton-x-100 for 1 hour at room temperature. Primary antibodies were applied overnight at 4°C, followed by washes and 1-hour incubation in secondary antibodies. Tissue sections were mounted in Prolong Gold with DAPI (Thermofisher) and imaged on a Zeiss Axioscan slide scanner and a Zeiss LSM 880 confocal microscope for high-resolution imaging.

| **Antigen target**<br>**Number** | **Working concentration** | **Vendor** | **Product** |
| --- | --- | --- | --- |
| Somatostatin (Sst) | 1:250 | Millipore | MAB354 |
| Somatostatin-14 (Sst) | 1:1000 | Penninsula | T-4103 |
| Parvalbumin (Pvalb) | 1:1000 | Sysy | 195004 |
| Vasoactive intestinal peptide (Vip) | 1:250 | Sysy | 443005 |
| Neuronal nitric oxide synthase (Nos1)<br>ab1376 |  | 1:500 | Abcam |
| Ctip2 | 1:500 | Abcam | ab18465 |
| Satb2 | 1:500 | Sysy | 327004 |
| GFP (reactive against EYFP) | 1:500 | Invitrogen | A-111222 |
| RFP (reactive against mCherry) | 1:250 | Chromotek | 5f8 |

### Tissue clearing and light sheet imaging

Animals were prepared as described above for morphological reconstruction. Animals were then transcardially perfused with ice cold 1xPBS followed by 4% PFA. Brains were then dissected and post fixed in 4% PFA at 4°C for 24hr. Brains were processed with SHIELD reagents (Life Canvas) before beginning active tissue clearing using the SmartBatch+ system (Life Canvas) using manufacturer protocols. Tissues were RI matched using EasyIndex (Life Canvas) and imaged on SmartSPIM light sheet microscope (Life Canvas) with a 3.9x magnification objective (NA: 0.2, Thorlabs manufactured modified by Life Canvas). Data aligned to the Allen CCF using NeuroInfo software (MBF Biosciences).

### Whole brain sparse labeling and reconstructions

Whole-neuron morphology (WNM) data were generated using HortaCloud ^94^, an open-source, cloud-based neuron reconstruction platform. Morphological reconstructions were performed on gene-defined, sparsely labeled individual neurons imaged across the entire mouse brain using two-photon fluorescent Micro-Optical Sectioning Tomography (fMOST) ^59,60^. For each neuron, the full local and long-range axonal arbor and dendritic trees were manually identified and traced by an experienced annotator. A second independent annotator then proofread the initial reconstruction adding missed branches and removing incorrectly identified segments. All branch points and terminal endings were verified to ensure complete reconstruction of each neurite. Any discrepancies between annotators were resolved through consensus before the reconstructed cell was deemed final.

### Slice electrophysiology and optogenetically evoked inhibitory postsynaptic currents (oIPSCs)

Acute coronal slices of the neocortex were prepared from adult Sst^flp^ ^+/+^ ; Nos1^creER^ ^+/-^ mice of either sex injected with AAV CreOn/FlpOn-ChR2-EYFP (1µL split across two depths) (DV -0.25 and DV -0.55) at AP-3.3, ML ±2.7, 5-eight weeks prior to recording. Mice were anesthetized with isoflurane and transcardially perfused with ice-cold NMDG-based cutting solution containing (in mM) 92 NMDG, 2.5 KCl, 1.25 NaH_2_PO_4_, 30 NaHCO_3_, 20 HEPES, 25 glucose, 2 thiourea, 5 Na-ascorbate, 3 Na-pyruvate, 0.5 CaCl_2_·2H_2_O, and 10 MgSO_4_·7H_2_O, adjusted to pH to 7.3–7.4 with 5 M hydrochloric acid. The brain was rapidly dissected and sectioned at 250-260µm a vibratome (VT1200S, Leica) and allowed to recover in the same solution for 5-10 min at 34°C before transfer to warm artificial cerebrospinal fluid (ACSF) containing (in mM) 120 NaCl, 25 NaHCO_3_, 1.25 NaH_2_PO_4_, 3 KCl, 1 MgCl_2_, 1.5 CaCl_2_, and 11 glucose, 3 Na-pyruvate, 1 Na-ascorbate equilibrated with O_2_ and 5% CO_2._ Slices were incubated for another 30min and then kept at room temperature until recording for up to eight hours. For a subset of experiments, we used a sucrose-based cutting solution containing (in mM) 90 sucrose, 60 NaCl, 5 MgCl_2_, 2.75 KCl, 1.25 NaH_2_PO_4_, 1.1 CaCl_2_, 9 Glucose, 26.5 NaHCO_3_, 3 Na-pyruvate, 1 Na-ascorbate, equilibrated with 95% O_2_ and 5% CO_2_. Slices were transferred directly to warm ACSF after cutting under these conditions.

We verified the presence of ChR2-EYFP-expressing fibers in each slice at various distances from the injection site prior to recording. Putative postsynaptic neurons were recorded across all cortical layers and at various distances from the injection site at near-physiological temperature ∼32 °C with an internal solution containing (in mM) 15 CsCl, 120 CsGluconate, 8 NaCl, 10 HEPES, 2 MgATP, 0.3 NaGTP, 0.2 EGTA, and 2 mg/ml biocytin for post hoc anatomical analysis (pH adjusted to 7.2 with CsOH; osmolarity adjusted to 290 mOsm). The chloride reversal potential for was ∼ -44 mV, and GABAergic currents were recorded at holding potential of 0 mV. Visually guided whole-cell recordings were obtained with patch pipettes of ∼3 MΩ resistance pulled from borosilicate capillary glass (BF150-86-10, Sutter Instrument, Novato, CA). Electrophysiology data was acquired using a Sutter dPatch amplifier (Sutter Instruments), digitized at 10 kHz and filtered at 5 kHz. oIPSCs were evoked with 0.5 msec long whole-field blue light pulses (CoolLED) at a frequency of 0.2 Hz. To isolate inhibitory currents in voltage-clamp the following receptor antagonists were added to the bath solution (in µM): 2 R-CPP, 5 NBQX, 1.5 CGP to block NMDA, AMPA and GABAB receptors, respectively. To confirm GABAergic identity oIPSCs were blocked with 50 µM gabazine SR95531 in some experiments. To confirm that oIPSCs were monosynaptic, 0.5 µM TTX and 100 µM 4AP were included in the bath solution in a subset of experiments. All drugs were purchased from Abcam (Cambridge, MA), Tocris (Bristol, UK) or HelloBio (Princeton, NJ). After recording, the patch pipette was withdrawn slowly to allow resealing of the membrane and slices were fixed in 4% PFA overnight. Biocytin-filled neurons were labeled with streptavidin-Alexa 647 using standard protocols. Confocal z-stacks of streptavidin-labeled neurons and EYFP+ Sst-Chodl fibers were taken on a Zeiss LSM 880 microscope at 1-2 µm increments. Z-stacks were processed in ImageJ and their position in the cortex recorded.

### Habituation and head restraint

Animals were briefly habituated to handling for several days prior to surgery. Animals were allowed to recover for 2 days before continuing habituation to handling. After a minimum of one week post-implantation, animals were gradually exposed to the head fixation apparatus (https://doi.org/10.25378/janelia.24691311), which consists of a low-friction rodent-driven belt treadmill. The headfixation duration was gradually increased over 2 weeks until animals were comfortable with multi-hour headfixation sessions.

To promote sleep while headfixed, animals were given multiple habituation sessions on the recording rig before data collection. We ensured that the treadmill was oriented horizontally with no slope, with the distance between the head fixation level and treadmill properly adjusted. The set up was thoroughly cleaned to minimize odors from other animals, and a radiant heat lamp was provided.

### Videography

Video of mice was acquired at 30Hz using Blackfly machine-vision ethernet-enabled cameras (FLIR) equipped with a Basler lens (model c125-1620-5m) for headfixed experiments and a Computar lens (TG4Z2813FCS-IR 0.33-Inch) for freely moving experiments. Rigs were illuminated with infrared LED arrays, similar to methods described previously ^95^. The camera was configured to send TTL pulses to synchronize videography with other data streams.

### In vivo 2 photon calcium imaging

GCaMP was expressed using either an AAV-CreOn/FlpOn-GCaMP6m viral vector or via transgenic expression in the Ai210 mouse line (a Cre and Flp dependent GCaMP7f). No substantial differences were observed between the two approaches. Animals were imaged on a custom Bergamo 2 photon microscope coupled to a Ti:Sapphire laser (Mira 700, Coherent). Emitted light was collected though a 10x 0.5 NA long-working-distance objective (TL10X-2P, ThorLabs). Images were typically acquired with ThorImage software at 1.4 frames/second with a resolution of 512x512 pixels and for experiments coupled with flexible probes, at 14.6 frames/second with a resolution of 128x128 pixels. For microprism data, cell positions were estimated based on the distance of the soma relative to the visible damage layer produced by the microprism insertion, which was reference as cortical surface (estimation relative to Allen Brain Atlas). Data streams were synchronized using TTL pulses collected on an RHD USB Interface Board (Intan Technologies) at 20kHz. Prior to imaging, novel objects were placed in the animal’s home cage, one per hour for four hours, to promote exploration and subsequently facilitate sleep during imaging.

### Acute in vivo electrophysiology and optogenetics

Animals were injected with AAV CreOn/FlpOn-ChR2-EYFP and implanted with headposts as described above. After viral expression, tamoxifen induction, and treadmill habituation, animals were administered dexamethasone (4mg/kg, i.p.) 1h prior craniotomy. A ∼3mm craniotomy was made over the injection site (V1).

A 64-channel linear silicon probe (H3 probe, Cambridge Neurotech) physically coupled to a tapered optical fiber was slowly (1µm/sec, over ∼20 minutes) inserted into V1. The craniotomy was kept moisturized during the recording by placing a small volume of silicon oil on the surface of the brain. State measurements were made as described below. Between recording days, the craniotomy was protected with Kwik-cast silicon elastomer (World Precision Instruments). Mice were recorded once per day for 3-4 days.

Recordings were split into blocks of ∼30 minutes of spontaneous periods (without stimulation) and ∼30-minute periods of stimulation. Several different types of stimulation were used including sinusoids (1Hz, 4Hz, 10Hz, 40Hz), flat pulses, and white noise, each delivered for 30-seconds followed by 30-seconds inter trial interval (ITI). Blue light was provided by a fibercoupled LED (MF470F4, Thorlabs). Signals to control LED light intensity were generated using custom written MATLAB software and delivered to NI-Daq card (NI-PCIe-6323) to be output to the LED control box (LEDD1B, Thorlabs). No light artifacts were detected in our recordings.

Across >700 recorded single units, plus additional multi-units from 10 animals, no optotagged units were found, underscoring the extremely low abundancy of this neuronal population and the difficulty of capturing them directly in silicon-probe recordings.

Data was acquired using an Intan RHD2000 interface board at 20kHz.

### Multi-LFPs and optogenetics

Animals were injected with AAV CreOn/FlpOn-ChR2-EYFP and implanted with headposts as described above. Recordings were split into blocks of ∼30minutes of spontaneous periods (without stimulation) and ∼30minute periods of stimulation using 10Hz sinusoid frequency. Similar to acute recordings, the stimulation periods consisted of 30 seconds of blue light followed by 30 seconds of inter trial interval (ITI). The same optical and acquisition hardware was used as described above.

### Freely moving behavioral assay

Animals were injected with AAV nEF-CreOn/FlpOn-hM3Dq-mCherry or control EF1a-fDIO-mCherry, implanted with LFP wires as described above and single-housed post-surgery. Mice were habituated to wired tethering via 16-channel digital headstage with accelerometer (Intan Technologies) and connected to a motorized commutator (Doric Lenses or Neurotek). To minimize stress and noise distraction, animals were kept in their home cage, inside an acoustic foam box for 2 hours per day for 3-5 days. Mice were also habituated to receiving systemic saline injections at the beginning of the session for at least 2 days before monitoring their behavioral state. A consistent time of the day was used for each mouse during the light/inactive phase (ZT3 to ZT7) or during the dark/active phase (ZT18), and maintained throughout the experiment. No differences in sleep behavior were detected across specific ZT windows used within the light/inactive phase, but sessions during light/inactive and dark/active phase were analyzed separately.

Following habituation, animals received i.p. injection of either Vehicle (saline) or Clozapine N-oxide (CNO, Hello Bio) at 0.5mg/kg, a dose shown not to impact sleep of control animals lacking DREADD ^96,97^ and confirmed in our control experiment (12 sessions from 6 animals during the light phase, Supp Fig 15). The initial treatment was counterbalanced across animals, and the alternate treatment was administered the following day. Thus, each session was paired with its opposite treatment for analysis (Vehicle *vs* CNO or CNO *vs* Vehicle). Animals completed two Vehicle and two CNO sessions, and data for each treatment was averaged across the two sessions. To ensure sufficient wakefulness, only animals that remained awake for at least 60% of the time during Vehicle sessions were included in the analysis within light/dark phase. We obtained a total of 28 sessions during the light phase from 14 animals and 12 sessions during the dark phase from 6 animals.

LFP data was acquired at 1250kHz using an Intan RHD2000 interface board.

### Behavioral state scoring

Behavioral state was divided based on a combination of the acquired state measurements. First, to score sleep states we utilized previously published and validated methods implemented in the Buzcode toolbox from the Buzsaki lab ^61,62^. In brief, this method provides automatic state scoring by distinguishing slow wave sleep (SWS) from wake using the slope of the power spectrum (a large slope occurs during high delta periods of sleep). Rapid eye movement (REM) sleep is identified as periods of high theta/delta ratio with low electromyogram (EMG). The automatic scoring is then visualized and manually refined by experts. Importantly, for optogenetic and chemogenetic experiments state was scored using LFPs recorded from outside the neocortex, in hippocampus, to provide an accurate measure of state.

The remaining periods of wakefulness were then divided into movement and quiet wake states. In headfixed recordings, movement intervals were defined as periods of ≥5sec in which facial motion energy (extracted using Facemap, binned at 1sec) exceeded the 90th percentile of the session’s distribution. Quiet wake was defined as the remaining wake intervals, with a minimum duration of 5 seconds. A similar procedure was applied in freely moving experiments, but accelerometer signals were used instead of the facial motion to classify movement and quiet wake states. We noted that headfixed mice exhibited substantially longer quiet-wake periods compared to freely moving animals, and we confirm, consistent with prior reports, that head-fixed mice sleep with eyes open, and display typical LFP signatures (i.e., high delta power during SWS and high theta/delta ratio during REM) ^98^. Although episodes of quiet wake, REM and SWS in headfixed mice may differ in meaningful ways from those recorded in in freely sleeping animals, we used the same state terminology across head-fixed and freely moving experiments for consistency.

### Data analysis

Data were analyzed using open-source software packages and custom written MATLAB and python code.

### Whole neuron morphology projection matrix

SWC files were merged and resampled so that spacing between nodes was uniform. Merged files were registered to the average mouse brain template of the Common Coordinate Framework (CCFv3, ^99^). CCF registered reconstructions were translated so that all somas were positioned on the left hemisphere. To generate a projection matrix, we took two strategies including using all axons present in an area and the other which required that a targeted structure contain at least one branch tip and one node. All projections were ipsilateral, and targeted structures are reported only for the ipsilateral hemisphere.

### Whole brain fiber density and rabies

Mice were cryosectioned as described above, with uniform sampling throughout the entire brain. Imaged brain sections were aligned to the Allen Common Coordinate Framework using Neuroinfo software (MBF Bioscience) using the *Brainmaker* workflow. Labeled neuronal arbor in aligned tissue was then reconstructed within single sections using Neurolucida 360 (MBF Bioscience). Density measurements for individual areas were made by calculating the total path length of reconstructed arbor within that area divided by the volume of the area (the 2D area within section multiplied by section thickness). The coordinates of each identified rabies-positive neuron were compiled to yield a complete whole-brain map for each mouse. To account for inter-animal variability in labeling efficiency, data from each brain were normalized to the total number of rabies-positive cells identified in that animal. These normalized values were then used for quantification of layer distributions, regional localization, and subregional visual cortex analyses.

### Ex vivo patch-clamp

Electrophysiology data were analyzed using SutterPatch (written in IgorPro, Wavemetrics). Optogenetically evoked inhibitory postsynaptic currents (oIPSCs) were aligned to the light stimulus and 20-60 traces were averaged to calculate oIPSC amplitude and latency. Amplitude was measured as the maximum positive peak of the baselined -subtracted oIPSC, and latency was calculated as the time between the light stimulus and crossing of a 2x SD threshold of the oIPSC from baseline.

### Extraction of calcium activity

Acquired images were processed using Suite2p ^100^ to extract ROIs and fluorescence traces. DeltaF/F0 values were computed by normalizing fluorescence to a moving 10th percentile within 10-min windows as the baseline (F0). Deconvolved calcium signals were obtained from suite2p, and due to their higher temporal resolution, were used to examine the temporal relationships between calcium activity, delta power, state transition and DOWN states. To investigate state transition, calcium activity was evaluated only during states lasting a minimum of 10 seconds. The latency of calcium activation during SWS was defined as the earliest time point at which the deconvolved calcium signal exceeded 2 standard deviations (SD) above baseline activity. When measuring calcium activity around DOWN states, calcium traces were corrected by subtracting chance-level calcium activity generated by circularly shifting the calcium signal (circshift in Matlab) 100 times within the 1 second window around the onset of DOWN states.

### Arousal score

Arousal score (Fig. 2b) was calculated using the *pca* function in Matlab. It corresponds to the time varying score for the first principal component of the measured state metrics. Principal component loadings calculated for each individual recording to provide robustness against measurement error for any singular state metric.

### Classification of low arousal vs high arousal correlated cells

Cell types were determined in Sst^flp^ ; Nos1^creER^ animals based on the correlation of deltaF/F traces with state metrics. K-means clustering (n=2) was performed on these correlation coefficients. Separately, we calculated the correlation coefficient of the deltaF/F signal with the arousal score. Cells with negative correlation coefficients (low arousal correlated) overlapped completely with the k-means low arousal cluster.

### Division of high vs low Sst-Chodl cell activity epochs

DeltaF/F traces were averaged within each recording and then z-scored. High activity epochs were defined as periods where activity was above 2 SD from the mean and low activity was below 0.5 SD.

### Single Units and LFP extraction

Single units were isolated using Kilosort2 ^101^. Figures contain data from both broad-spiking and narrow-spiking units as no differences were found between these groups. LFP signals were collected from the microwire array or from silicon probes and were downsampled to 1250 Hz. EMG signals were obtained by band-pass filtering the raw signal (>200 Hz) followed by a Hilbert transform.

### Calculation of power spectra

Power spectra were calculated using a wavelet-based spectrogram method (Morlet wavelets, 5-cycle width). These wavelets were generated for 100 log-spaced frequencies from 1Hz to 128Hz. For freely moving optogenetic experiments, changes in power spectra was assessed by comparing the 30 seconds of stimulation periods to the 30 seconds of inter trial interval (ITI). To investigate changes in power spectra across the neocortex as a function of distance from the injection site, we selected, at each distance (Proximal to Anterior), the channel showing the largest delta-band power increase. This approach was implemented to minimize the confounding effects of asymmetric projections where Sst-Chodl fibers might be absent on one side but present on the other at a single distance, given the substantial in individual projection patterns (see Figure 1).

### Determination of cortical depth

As shown previously, high frequency power (>500 Hz) peaks in mid layer 5 ^102^. We interpolated between this layer 5 channel and the first channel outside of the brain (1.3 mm probe inserted 1.2 mm into the brain) assuming the remaining distance, without the mechanical deformation of the brain caused by insertion of the probe, to be 700µm. With these interpolated pseudo-depths, we then defined layers boundaries at 100, 300, 450, 650, and 900µm.

### Spiking synchrony

The cross correlation was calculated for each unit pair using default parameters of the CCG.m function from Buzcode package. The spike times of these units were shuffled, and the cross correlation was recalculated 100 times to estimate a chance level. The real cross correlation was then normalized by this chance level to quantify spiking co-activation.

### Phase locking

The Fourier spectrum of the LFP was calculated for each spike from each unit using the ft_spiketriggeredspectrum function from the fieldtrip toolbox. The circular mean of the phase values at each frequency was calculated as the phase locking value. Though this measure is sensitive to spike count, with higher spikes leading to higher phase locking values, we believe this is not an issue because stimulation leads to slightly lower firing rates compared to baseline.

### DOWN state detection

Down states were detected using a previously published method ^65^. Briefly, the method detects a confluence of a large positive deflection in the LFP, a drop in gamma band power, and a sharp drop in multi-unit firing rate.

Linear regression model

Deconvolved calcium signals were used to predict z-scored delta amplitude after constructing a time-lagged design matrix spanning a 20 second temporal window, sampled every 0.2 seconds. The kernel was centered on the present time point allowing both past and future lags to contribute to the prediction. Ordinary least-squares regression was fitted in a 5-fold cross-validation scheme, and model performance was quantified by the cross-validated coefficient of determination (R²). The resulting regression weights were averaged across folds to obtain a mean temporal kernel describing the influence of calcium activity at different lags on the predicted delta signal.

### Mouse tracking

Cage position and mouse nest area were manually delimited using a compilation of frames extracted every 10 min from each video. Animal position was detected from the body centroid of the mouse using DeepLabCut open-source system ^68^. The distance moved by the mice was extracted according to the registered position of the cage. The duration spent in the nest was estimated by calculating the time of the mouse present in the nest according to the total duration of the experiment.

### Allen Brain Observatory data analysis

Sst expressing cell state-correlation data was retrieved using the allensdk and querying the database for all experiments performed with Sst^cre^ and Cre-dependent GCaMP mice. Pearson’s correlations were calculated between activity traces and state metrics after binning both signals into 3 second windows. Data on proportions of Sst-Chodl cells with Sst population was retrieved from the dataset presented in ^32^ from the web-based Brain Knowledge Platform (https://knowledge.brain-map.org/data).

### Bugeon et al 2022 data analysis

Data was retrieved from https://doi.org/10.6084/m9.figshare.19448531; and analyzed from spontaneous recordings without visual stimulation.

The following software packages were used for analyzing the data presented in this paper:

| Package name | URL | Publication |
| --- | --- | --- |
| allensdk | https://github.com/AllenInstitute/AllenSDK | N/A |
| buzcode | https://github.com/buzsakilab/buzcode | N/A |
| facemap | https://github.com/MouseLand/facemap | Syeda et al 2023 |
| suite2p | https://github.com/MouseLand/suite2p | Pachitariu et al 2017 |
| kilosort2 | https://github.com/MouseLand/Kilosort | Pachitariu et al 2023 |
| fieldtrip | https://www.fieldtriptoolbox.org/ | Oostenveld et al 2011 |
| deeplabcut | https://github.com/DeepLabCut/DeepLabCut | Mathis et al 2018 |
| Phy2.0 beta | https://github.com/cortex-lab/phy | N/A |

### Statistical analysis

Matlab (MathWorks) was used for statistical analysis. No power calculations were used to pre-determine sample sizes or to formally assess normality. Sample sizes were determined based on previous literature. Comparisons were performed using two-tailed parametric t-test (*ttest*), one-way, two-way (*anova1*), or repeated measures ANOVA, with post hoc Bonferroni corrections for multiple comparisons (unless stated otherwise). Differences in proportions were assessed using Chi-square tests (*chi2cdf*). For optogenetic experiments (linear probes, multi-LFPs and patch clamp) and comparison of SWS duration between light/dark sessions, linear mixed-effects model – lme, was implemented in MATLAB (fitlme) – and the model significance was assessed using ANOVA, which tests for the contribution of each fixed factor and their interaction while accounting for unequal sample sizes. Values and statistical tests used are reported in the text and data are represented as mean ± standard error mean (s.e.m.) unless reported otherwise. Significance was set with alpha = 0.05 and was represented on graphs as the following: *P < 0.05, **P < 0.01, ***P < 0.001.

## Supplementary Figures

**Supplemental Figure 1.**
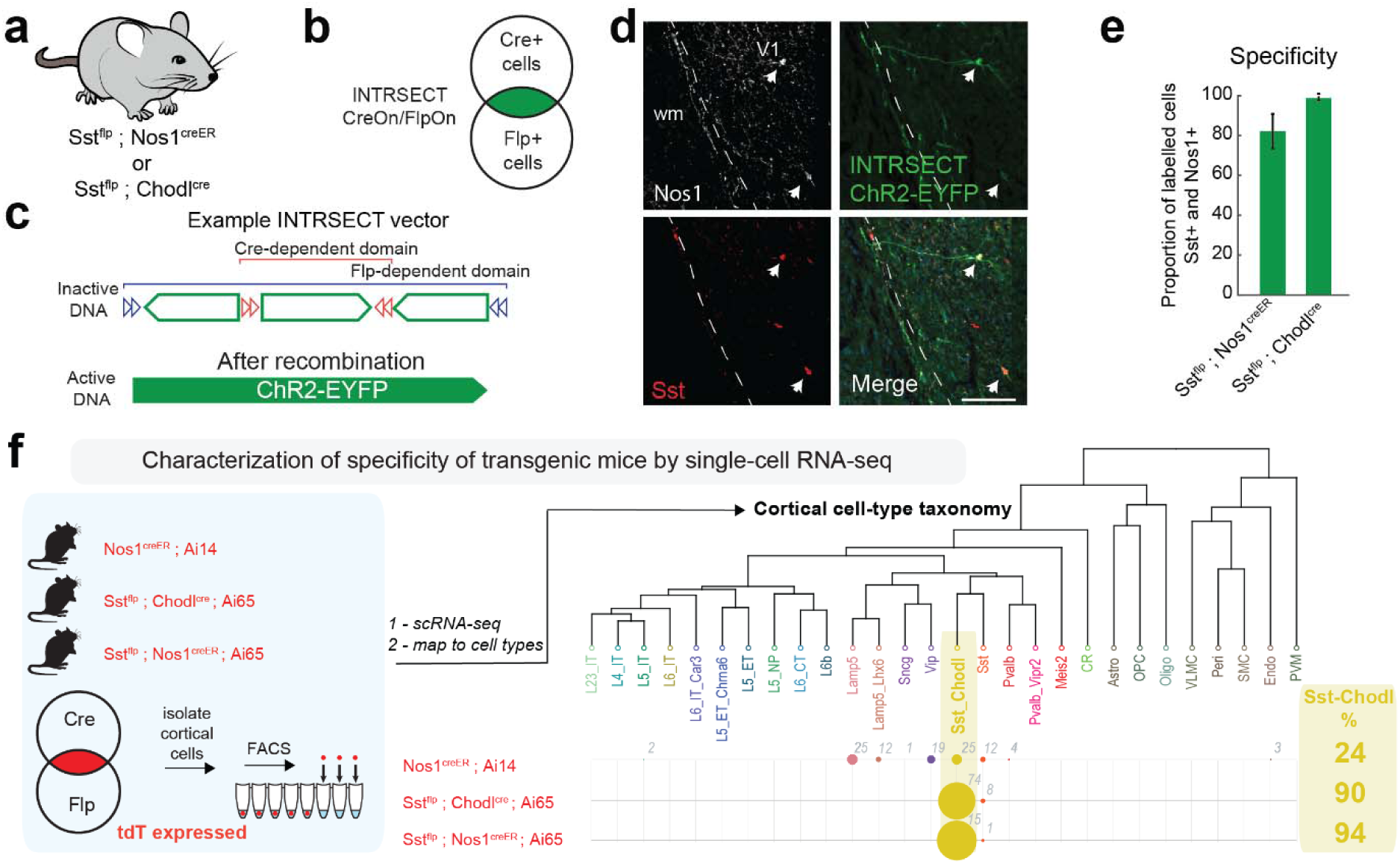
Intersectional strategies to target Sst-Chodl cells with high specificity. **a)** Mouse lines used for intersectional targeting of Sst-Chodl cells. **b)** Venn diagram showing the populations labeled by CreOn/FlpOn INTRSECT vector. **c)** Example INTRSECT construct illustrating that fluorophore expression occurs only after both Cre and Flp recombination. This ensures highly selective access to Sst-Chodl cells. **d)** Histology from Sst^flp^ ; Nos1^creER^ mice showing ChR2-EYFP expression in Sst-Chodl cells in V1; arrows indicate cells co-expressing Sst and Nos1. Long-range axonal segments travelling through white matter (wm) are visible (scale: 150µm;). These examples show the specificity of labeling and the presence of projecting axons originating from targeted cells. **e)** Proportion of cells expressing GCaMP, Sst, and Nos1 in Sst^flp^ ; Chodl^cre^ mice (62 cells from 4) mice and Sst^flp^ ; Chodl^cre^ mice (30 cells), showing the specificity of the labeling strategies for studying Sst-Chodl cells. **f)** Cortical cell-type taxonomy obtained from single-cell RNA-sequencing of labeled neurons from three transgenic lines (Nos1^creER^, Sst^flp^ ; Chodl^cre^ and Sst^flp^ ; Nos1^creER^) crossed with the tdTomato reporter line (data from ^90^). Numbers indicate cell count. This transcriptomic analysis further verifies that the intersectional strategies provide enrich labeling of Sst-Chodl cells.

**Supplemental Figure 2.**
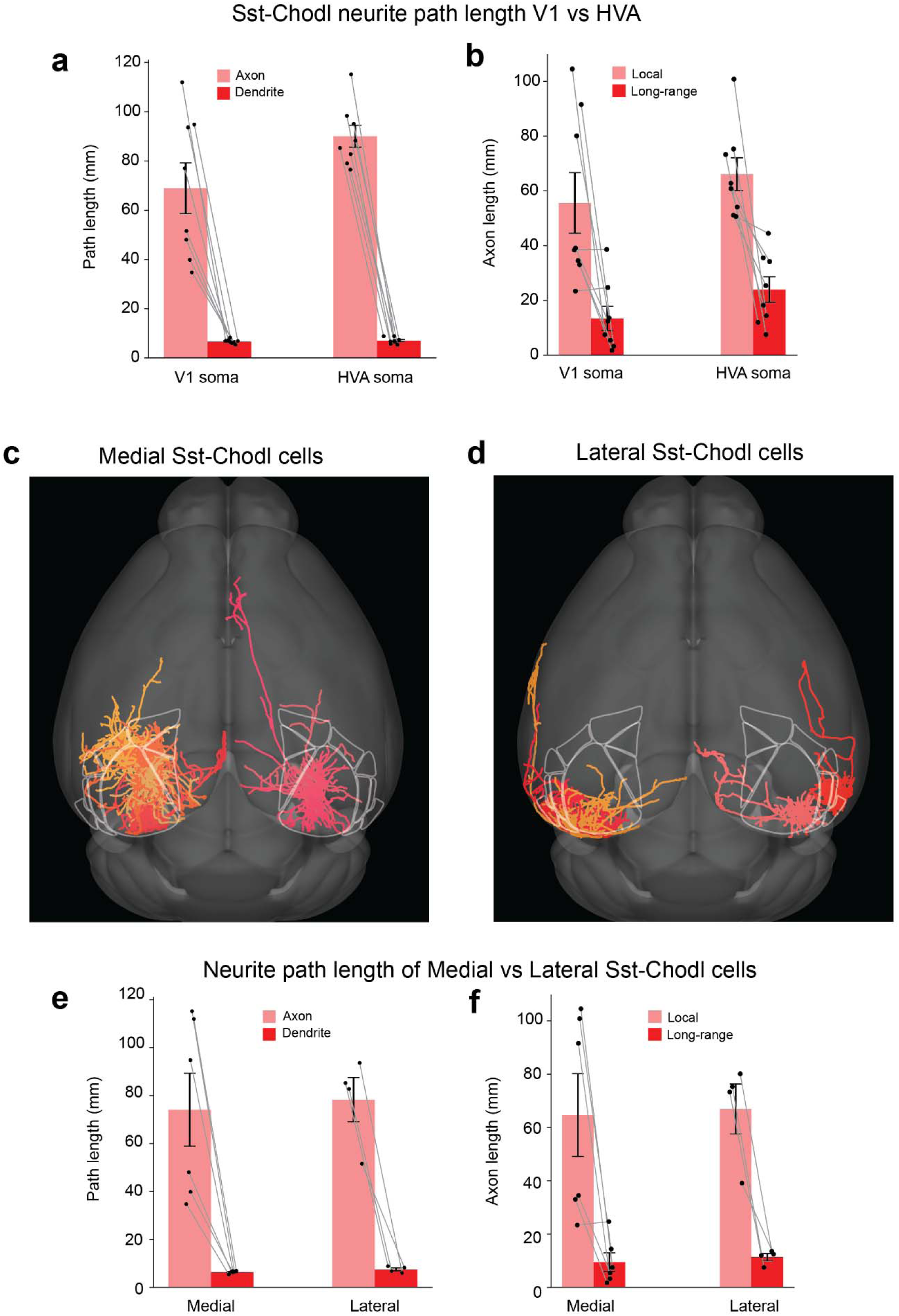
Reconstructed Sst-Chodl cells show consistent long-range projection patterns across cortical locations. **a)** Axonal and dendritic path length of reconstructed Sst-Chodl cells originating in primary visual cortex (V1) and higher visual areas (HVA). These measurements show that Sst-Chodl cells in V1 and HVAs have broadly similar arborization magnitudes. **b)** Same as (a), separated into local axon (within visual areas) versus long-range axon (outside visual areas). Both V1- and HVA-derived cells exhibit substantial long-range projections, indicating that long-range targeting is a shared property across visual cortical Sst-Chodl populations. **c-d)** Example reconstructions of Sst-Chodl cells with somata positioned in lateral vs medial regions of visual cortex. **e-f)** Quantification of total axonal and dendritic path lengths (e) and local versus long-range axon length (f) for medial vs. lateral Sst-Chodl cells. These comparisons show that cell-body location along the medial–lateral axis does not substantially alter overall arbor size or long-range projection extent.

**Supplemental Figure 3.**
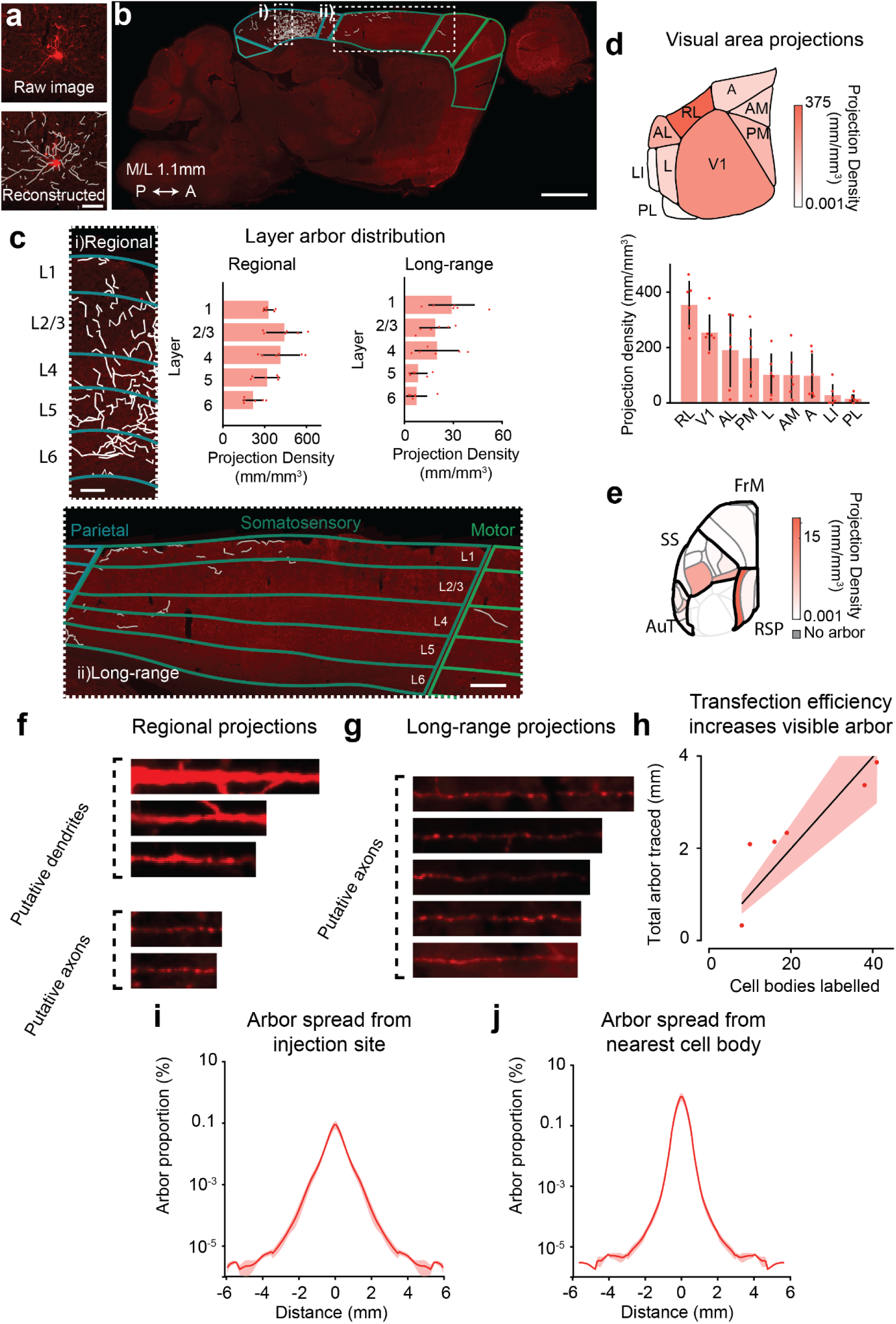
Dendritic and axonal arbors of Sst-Chodl cells reveal dense local innervation and widespread long-range projections. **a)** Example thin-section reconstruction of a Sst-Chodl cell arbor (scale: 100µm). **b)** Example thin-section showing both regional and long-range oScarlet-labeled axons from V1 Sst-Chodl cells (scale: 1mm). This reconstruction illustrates the coexistence of dense local arbors and extensive interareal projections. **c)** Insets from (b) with quantifying regional and interareal arbor density (scale: 100 and 200µm, respectively). These measurements show that visual areas receive the densest innervation, with substantial additional arbor distributed across other cortical regions. **d)** Projection density across visual cortical subregions averaged across mice, demonstrating consistent local arbor patterns across individuals. **e)** Quantification of long-range projections targeting neocortical areas outside the visual cortex, confirming broad ipsilateral interareal connectivity. **f)** Putative dendrites and axons near the injection site in V1. **g)** Putative axons lacking dendrites far from injection site (outside visual areas), consistent with distal long-range projections. **h)** Regression relating transfection strength to total arbor traced, indicating that measured arbor size scales with expression level. **i)** Spread of labeled arbor from the inferred injection site extending multiple millimeters. **j)** Spread of arbor from the nearest reconstructed soma, likewise travelling multiple millimeters, underscoring the long-range nature of Sst-Chodl projections. V1 = primary visual area, AL = anterolateral visual area, PM = posteromedial visual area, L = lateral visual area, AM = anteromedial visual area, A = anterior area, LI = laterointermediate area, PL = posterolateral visual area, AuT = auditory/temporal cortex, RSP = retrosplenial cortex, SS = somatosensory, FrM = frontomotor cortex. N = 6 mice. Data are means; shading and bars indicate s.e.m.

**Supplemental Figure 4.**
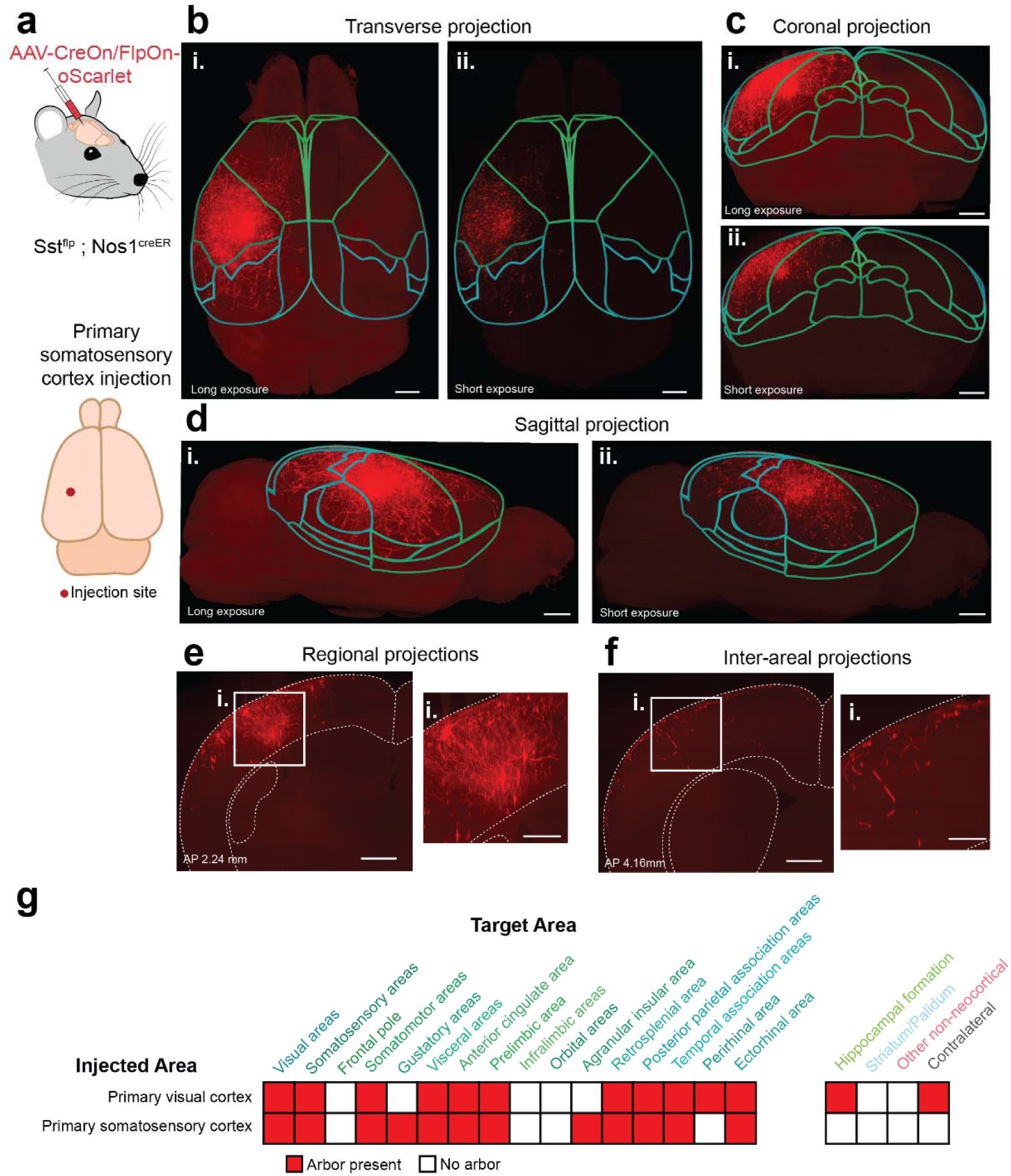
Sst-Chodl cells in the primary somatosensory cortex also send dense local arbors and widespread long-range cortical projections. **a)** Schematic of AAV-CreOn/FlpOn-oScarlet injection into primary somatosensory cortex (S1) of a Sst^flp^ ; Nos1^creER^ mouse to label S1 Sst-Chodl cells. **b)** Dorsal view of a cleared whole brain showing oScarlet-labeled S1 Sst-Chodl arbors with with (i) long and (ii) short exposure. **c-d)** Same dataset shown from anterior (c) and lateral (d) perspectives (scale for b–d: 1 mm). These views reveal dense local labeling in S1 and prominent long-range axons projecting across extensive ipsilateral cortical territory. **e)** Coronal section near the injection site showing dense local arborization (scale: 0.5 mm; inset 0.25 mm). **f)** Coronal section far from the injection site, where labeled fibers appear without nearby somata, demonstrating distal long-range projections from S1 Sst-Chodl cells. Insets correspond to represent zoomed regions noted in (i). **g)** Comparison of targeted cortical areas following injections into V1 vs S1, showing that Sst-Chodl cells originated from each region innervate surrounding neocortical regions but also send inter-areal projections.

**Supplementary Figure 5.**
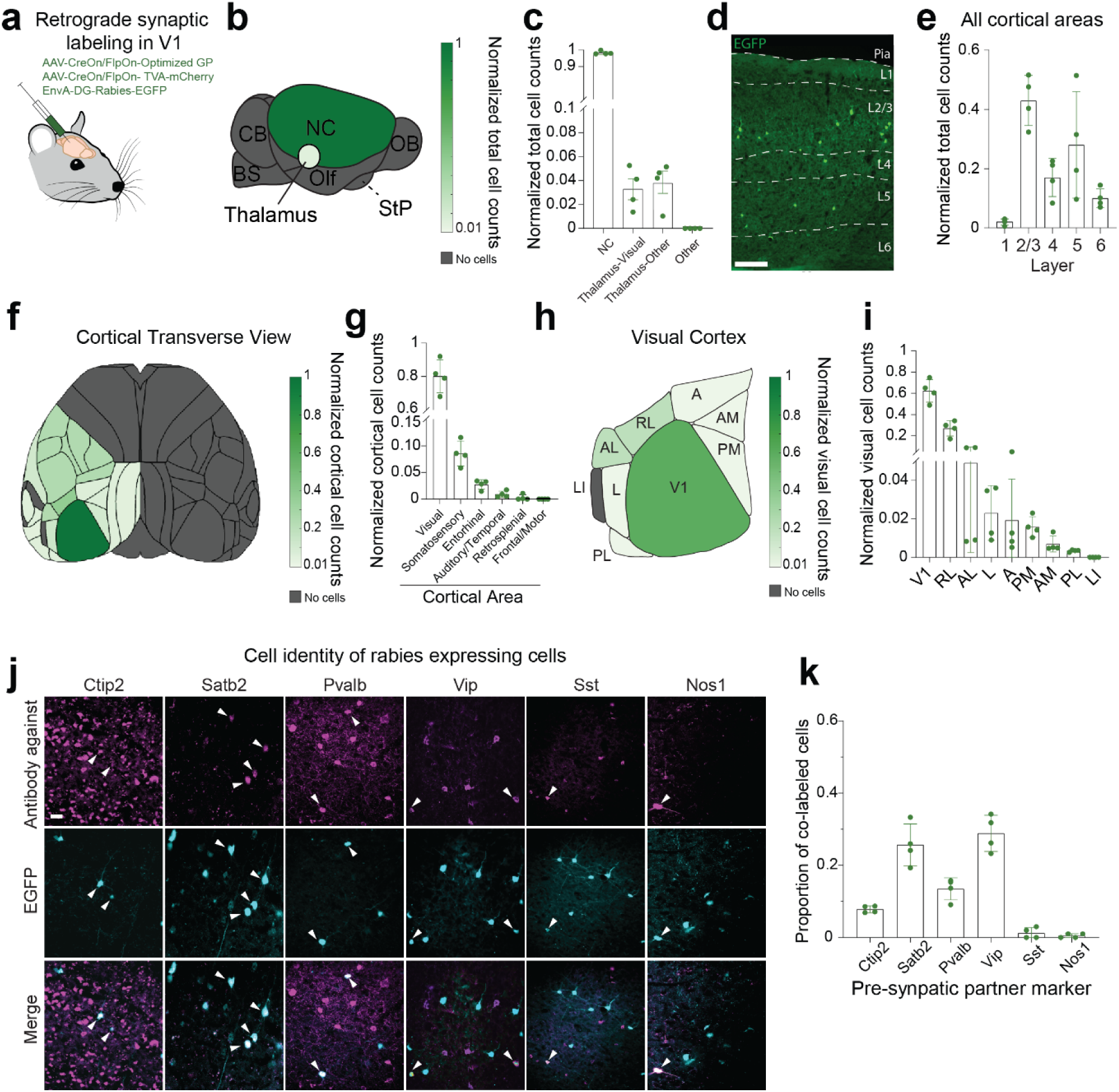
Sst-Chodl cells receive widespread intracortical inputs from diverse excitatory and inhibitory classes. **a)** Schematic of the monosynaptic rabies-based retrograde tracing strategy using intersectional genetics to identify presynaptic partners of Sst-Chodl cells. This approach enables selective labeling of neurons providing direct input to Sst-Chodl cells. **b)** Distribution of rabies-tagged pre-synaptic neurons across major brain structures. Only a small minority of labeled cells were found in the thalamus (28 out of 527 total cells), indicating that Sst-Chodl cells receive relatively sparse thalamic inputs. **c)** Cell counts across brain regions normalized to total labeled cells. “Thalamus-Visual” includes the dorsal lateral geniculate complex; “Thalamus-Other” includes dorsal thalamus and medial geniculate regions. **d)** Example coronal section showing rabies-expressing presynaptic neurons (EGFP) across layers of V1 (scale: 100 µm. **e)** Quantification of laminar distribution within the neocortex, expressed as a proportion of labeled neocortical cells. Inputs arise predominantly from intermediate layers (layers 2–5). **f-g)** Distribution and quantification of pre-synaptic neurons across neocortical areas, normalized to the total neocortical presynaptic pool (f, transverse view; g, quantification). Labeled neurons were almost exclusively ipsilateral and most abundant in V1. **h-i)** Distribution and quantification of presynaptic neurons across visual cortical areas, showing that V1 provides the densest inputs within the visual system (h, transverse view; i, quantification). **j)** Immunohistochemical examples of presynaptic cells co-expressing rabies and major neuronal subtype markers, including glutamatergic markers Ctip2 (putative pyramidal tract) and Satb2 (intratelencephalic), as well as GABAergic markers Pvalb, Vip, and Sst (scale: 20 µm; white arrows indicate co-labeled cells). **k)** Proportion of presynaptic neurons co-expressing each marker. These analyses show that Sst-Chodl cells receive input from all major cortical excitatory and inhibitory populations probed, with Satb2⁺ and Vip⁺ cells being the most prominent contributors and Sst⁺Nos1⁺ neurons the least frequent. Together, these data demonstrate that Sst-Chodl cells integrate a broad array of intracortical inputs, with minimal thalamic contribution and substantial excitatory and inhibitory presynaptic diversity. NC = neocortex; CB = cerebellum; NAc = nucleus accumbens; OB = olfactory bulb; BS = brainstem; Olf = olfactory areas; StP = striatum/pallidum; DLGN = dorsolateral geniculate nucleus; V1 = primary visual cortex; RL = rostrolateral visual area; AL = anterolateral visual area; L = lateral visual area; A = anterior area; PM = posteromedial visual area; AM = anteromedial visual area; PL = posterolateral visual area; LI = laterointermediate area. N = 4 mice. Data are means; bars indicate s.e.m.

**Supplemental Figure 6.**
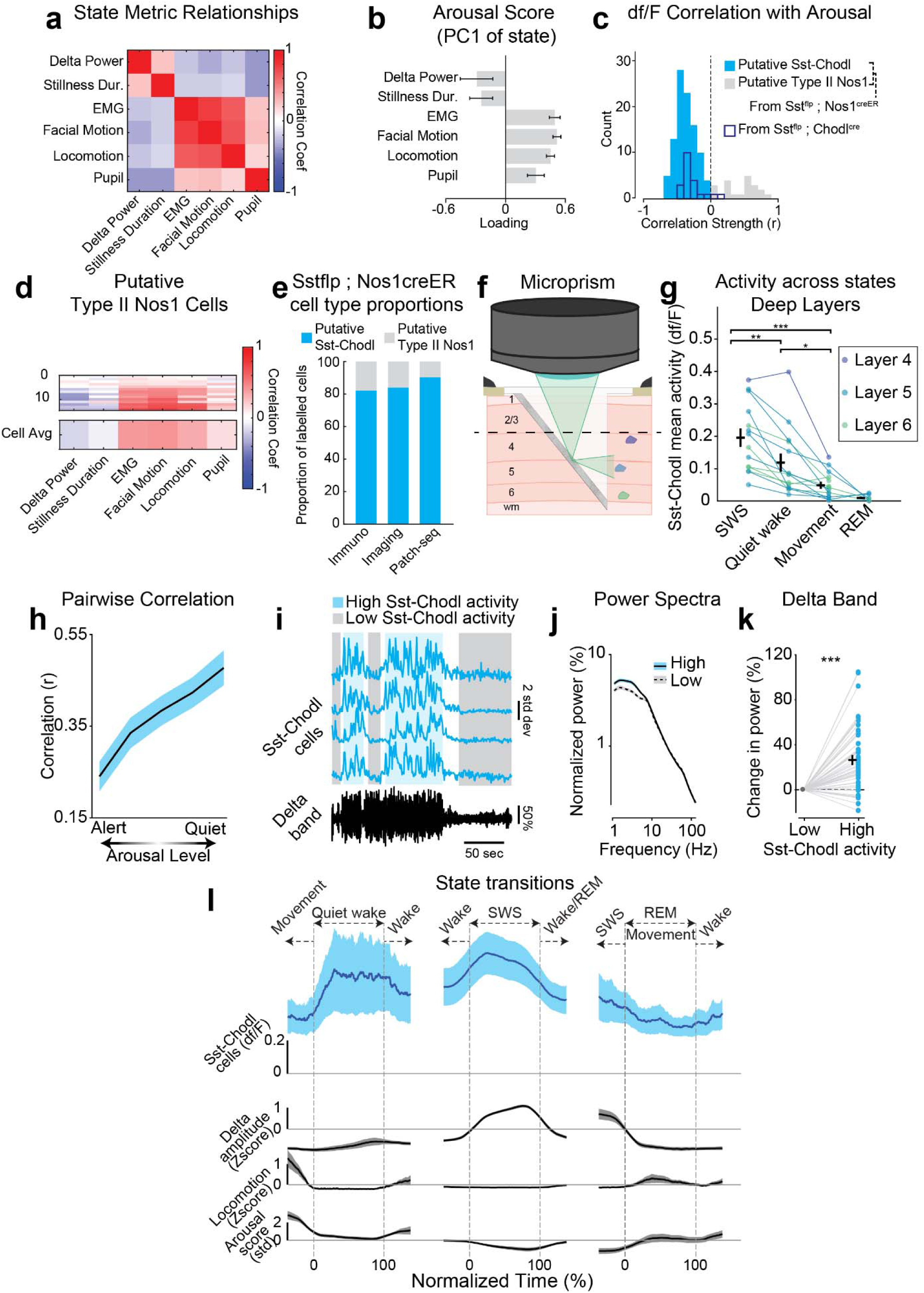
Population-level analysis shows that Sst-Chodl cells are most active during low-arousal, synchronized states. **a)** Average correlation between state measurements showing consistent relationship between the metrics. **b)** Consistent PC1 loadings in PCA of state measurements across animals. **c)** Distribution of correlation coefficients between imaged cell df/F and arousal score. Cells from Sst^flp^ ; Nos1^creER^ mice (full) and from Sst^flp^ ; Chodl^cre^ mice (clear) are assigned by k-means clustering for each space of df/F *vs* state metric correlation. **d)** Correlation of putative Type II Nos1 cell activity with state metrics (n = 16 out of 111 recorded cells from 15 animals from layers 1 to 3), showing a small proportion of cells with opposite pattern of activity as compared to putative Sst-Chodl cells. **e)** Proportion of Sst-Chodl and putative Type II Nos1 cells with identity determined by immunohistochemistry (Immuno, n = 62 cells from 4 animals), imaging and activity correlation (Imaging, n = 90 cells from 11 mice), and by patch-seq in Sst^flp^ ; Nos1^creER^ mice crossed with Ai65 mice from Allen Brain Institute in ^32^ (Patch-seq, n = 87 cells). The small proportion of putative Type II Nos1 cells is consistent across our immunohistochemistry and imaging results and sequencing data. **f)** Schematic of Sst-Chodl cells recorded in deep layers (estimated cell position in layer 4, dark blue; layer 5, light blue; layer 6, light green) with a microprism implanted in V1. **g)** Mean activity of deep layer recorded cells across states. ANOVA p < 0.001, post hoc multiple comparisons: SWS-QW ** p = 0.002, SWS-Movement *** p < 0.001, QW-Movement *, p = 0.016. N = 5 animals (from Sst^flp^ ; Chodl^cre^ mice x Ai210 cross), n = 14 cells. Putative Type II Nos1 cells were not represented in the graph (2 out of 16 recorded cells). Similarly to Sst-Chodl cells recorded in superficial layers, deep layer cells are mainly active during low arousal states, such as SWS and quiet wake. **h)** Mean pairwise correlation between cells based on arousal level (n = 42 pairs), showing consistent correlation activity across cells relative to arousal. **i)** Example recordings during periods of high and low Sst-Chodl cell activity. **j)** Power spectra during periods of high and low Sst-Chodl cell activity and **k)** quantification of delta band power change between these periods, p < 0.001 n = 97 cells, 15 animals. These results show that Sst-Chodl cells are most active when cortical networks are highly synchronized. **l)** Mean Sst-Chodl cell activity, delta amplitude, locomotion, and arousal level (note multiple scales) around state transitions. Sst-Chodl cell activity increases as animals enter low arousal states in parallel with rising delta power and decreases when they transition to desynchronized states. Data from this panel were obtained from 2photon recordings with flexible probes as presented in main Figure 2 e-k), n = 17 cells, 4 animals. ***: p < 0.001, **: p < 0.01, *: p < 0.05. Data are means; shading and bars indicate s.e.m.

**Supplemental Figure 7.**
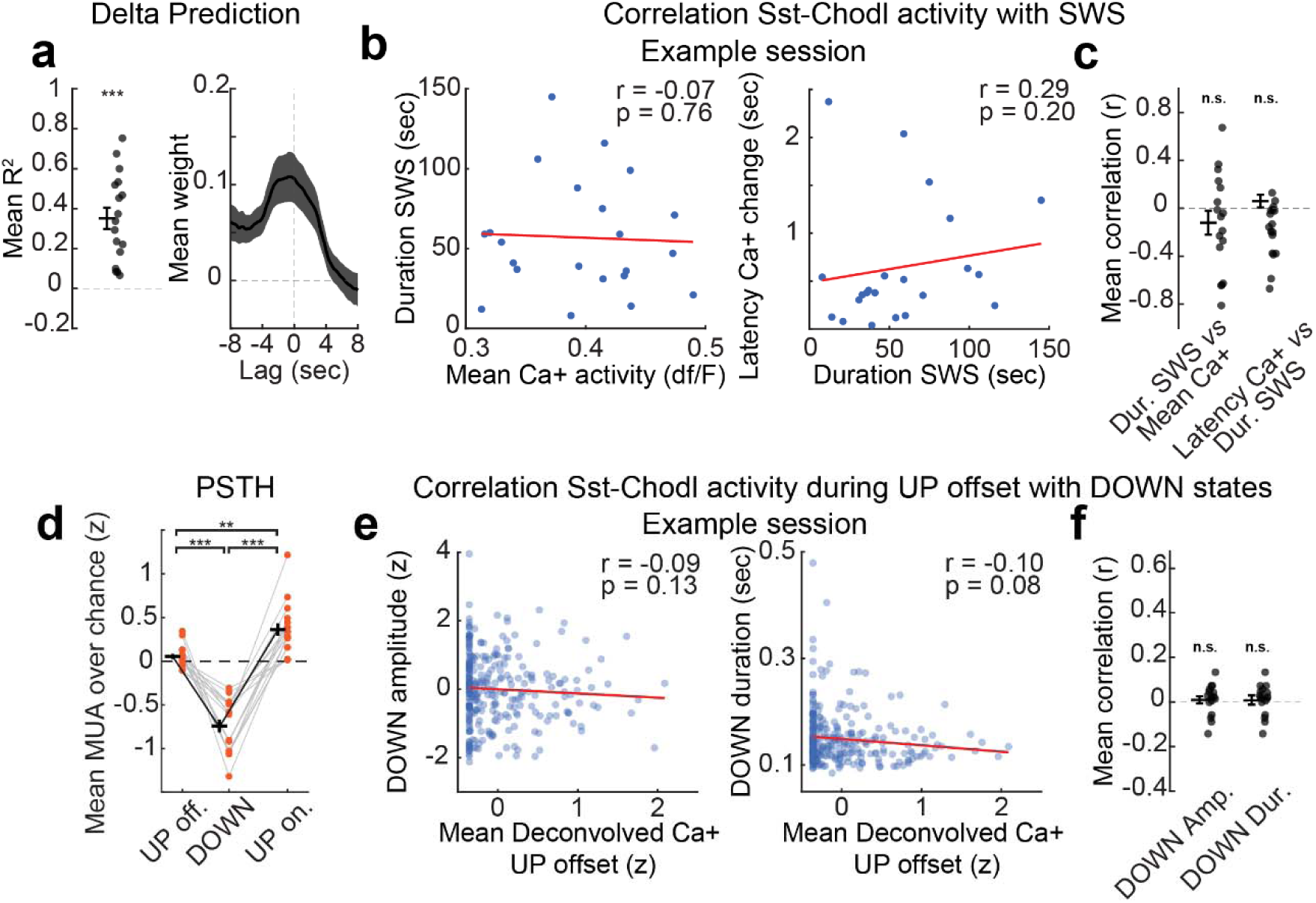
Sst-Chodl cell activity reflects ongoing SWS dynamics but does not predict SWS bout features. **a)** Average linear regression model predicting delta amplitude from deconvolved Sst-Chodl calcium activity. Left: cross-validated R² from ordinary least-squares regression (p < 0.001, one-sided Wilcoxon test). Right: mean kernel weights (±1 SD) across temporal lags. **b-c)** Relationship between SWS bout duration and Sst-Chodl cell activity. **b)** Example session showing correlations between bout duration and mean calcium activity (left: r = -0.07, p = 0.76) and between bout duration and activation latency (right: r = 0.29, p = 0.20; Spearman correlation). **c)** Mean correlation coefficients across cells (Duration SWS vs. Mean Ca^2+^, p = 0.266; Latency Ca+ vs. Duration SWS: p = 0.237; one-sided Wilcoxon test). These results indicate that Sst-Chodl activity does not predict the duration or onset timing of SWS bouts. **d)** Average PSTH of multi-unit activity (MUA) aligned to DOWN states: 400 msec before DOWN states (UP offset), during DOWN states, and 400 msec after DOWN states (UP onset). MUA shows a strong decrease at DOWN onset and rebound at UP onset (ANOVA p < 0.001; UP off.-DOWN p < 0.001; DOWN-UP on. p < 0.001; UP off.-UP on. p = 0.006). **e-f)** Relationship between DOWN-state features and Sst-Chodl cell calcium activity. **e)** Example session showing correlations between DOWN amplitude and mean UP-offset calcium activity (left: r = -0.09, p = 0.13) and between DOWN duration and mean UP-offset calcium activity (right: r = -0.10, p = 0.08; Spearman). **f)** Mean correlation coefficients across cells (DOWN amplitude vs. Ca^2+^: p = 0.332; DOWN duration vs. Ca^2+^: p = 0.586; Wilcoxon). These analyses show that although Sst-Chodl cells increase activity prior to DOWN onset, their activity does not predict the amplitude or duration of the upcoming DOWN state. Data in all panels are from simultaneous 2-photon and flexible-probe recordings (main Fig 2e-k), n = 17 cells, 4 mice. ***: p < 0.001, **: p < 0.01, ns, not significant. Data are means; shading and bars indicate s.e.m.

**Supplemental Figure 8.**
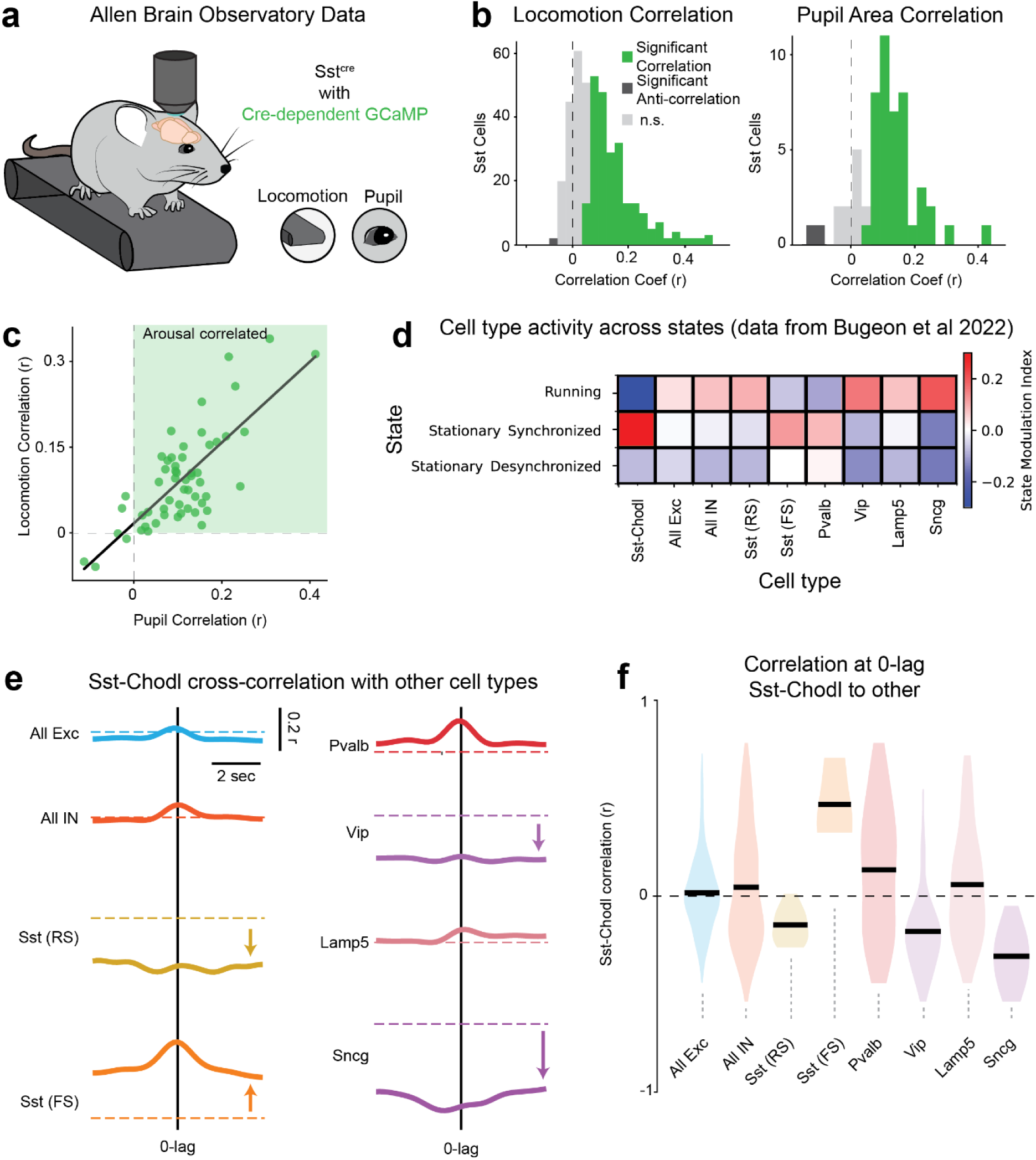
Sst-Chodl cells differ from other Sst and inhibitory subtypes in their modulation by arousal state. **a)** Schematic of Allen Brain Observatory recordings in which Sst cells are imaged with 2-photon calcium imaging while pupil diameter and locomotion are monitored. **b)** Distribution of correlations between Sst cell activity and locomotion (left; n = 404 cells from 47 sessions) and pupil diameter (right; n = 55 cells from 9 sessions). Most Sst cells show positive correlations with arousal-linked measures. **c)** Scatterplot of pupil vs locomotion correlations for Sst cells (n = 55 cells from 9 sessions), illustrating that Sst cells in these datasets predominantly increase activity with heightened arousal. **d)** Correlations between activity and arousal state across major neuronal subtypes (dataset from ref. ^103^). Cell counts: 6684 Exc, 1954 IN, 414 Pvalb, 87 Sst (regular-spiking; RS), 24 Sst (fast-spiking; FS), 2 Sst-Chodl, 392 Vip, 543 Lamp5, 53 Sncg cells. This comparison shows that most inhibitory and excitatory classes are positively modulated by arousal, whereas Sst-Chodl cells exhibit the opposite pattern. **e)** Cross-correlation between Sst-Chodl cells and other neuronal subtypes, showing their relationships to arousal-modulated populations. **f)** Distribution of zero-lag correlations between Sst-Chodl cells and other cell types. Cell counts: 812 Exc, 368 IN, 56 Pvalb, 8 Sst (RS), 3 Sst (FS), 51 Vip, 77 Lamp5, 10 Sncg. Fast-spiking Sst cells include Sst-Tac1-Tacr3 and Sst-Tac1-Htr1d subtypes; regular-spiking Sst cells include all non-FS and non-Chodl Sst subtypes. Black lines denote means. These datasets show that Sst-Chodl cells are negatively correlated with arousal-activated inhibitory and excitatory populations and most positively correlated with fast-spiking Sst subtypes.

**Supplemental Figure 9.**
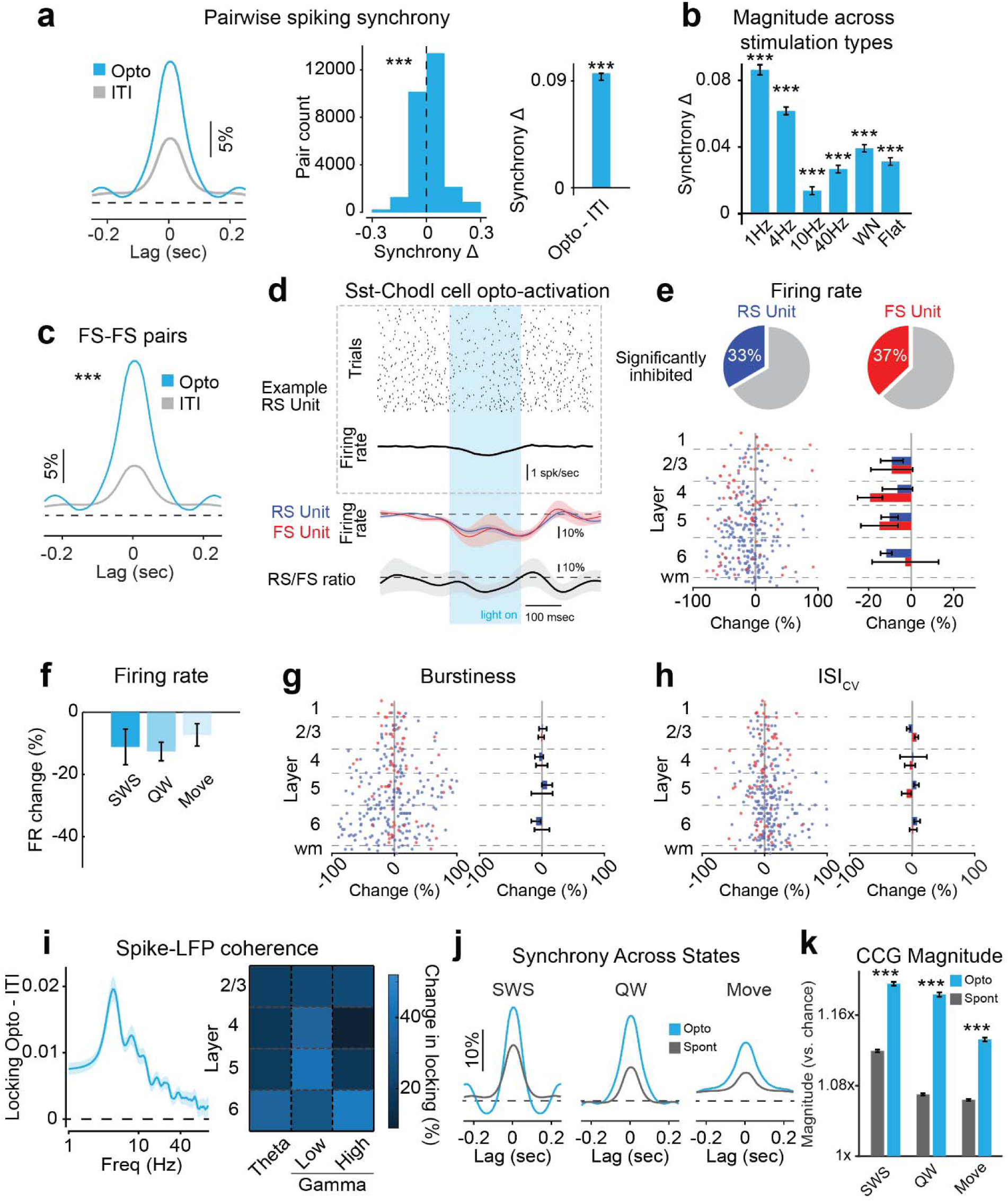
Optogenetic activation of Sst-Chodl cells with optogenetics increases spiking synchrony across neuronal subtypes and cortical layers. **a)** Normalized cross-correlograms (CCGs) comparing inter-trial intervals (ITIs) and stimulation periods. Distribution (left) and average (right) of CCG peak changes for all neuron pairs (p < 0.001, paired t-test). These analyses show that stimulation reliably increases pairwise spiking synchrony. **b)** Change in spiking synchrony across different stimulation protocols (p < 0.001, paired t-test; Opto vs. Spont), demonstrating that synchrony enhancement is robust across stimulation types. **c)** Spike-spike synchrony changes for fast-spiking (FS) cell pairs during Sst-Chodl cell stimulation (p < 0.001, paired t-test; n = 19,918 pairs) revealing that FS interneurons also exhibit increased synchrony. **d)** Top: example regular-spiking (RS) unit response to Sst-Chodl cell stimulation. Bottom: average responses of RS and FS units to stimulation and RS:FS response ratio, indicating minor firing-rate modulation relative to synchrony changes. **e)** Top: proportion of significantly modulated RS and FS units (paired t-test, pre-stim vs. post-stim). Bottom: firing-rate changes as a function of cortical depth (Opto vs. ITI). These results show small but systematic rate reductions in some units. **f)** Firing-rate changes across behavioral states (Opto vs. ITI) confirming that synchrony increases cannot be explained by increased firing activity. **g)** Change in burstiness (autocorrelogram ratio 1.5-13/ 200-300 msec) across layers (Opto vs. Spont). **h)** Change in coefficient of variation of interspike intervals across layers (Opto vs. Spont). Together, these metrics indicate minimal changes in spike structure despite increased synchrony. **i)** Spike-LFP coherence across frequency bands and layers, showing significant increases during stimulation (Stimulation p < 0.001; no significant effects of Layer or Frequency band; linear-mixed effects model). **j**) Change in spiking synchrony across behavioral states (all cells, Opto vs. Spont). **k)** Change in CCG peak synchrony across states (± 50 msec window; p<0.001 for all, paired t-test), indicating robust increases regardless of state. These experiments were performed in V1 of head-fixed mice. N = 14 sessions across 10 mice; n = 758 units and 32,710 pairs. ***: p < 0.001. Data are means; shading and bars indicate s.e.m.

**Supplemental Figure 10.**
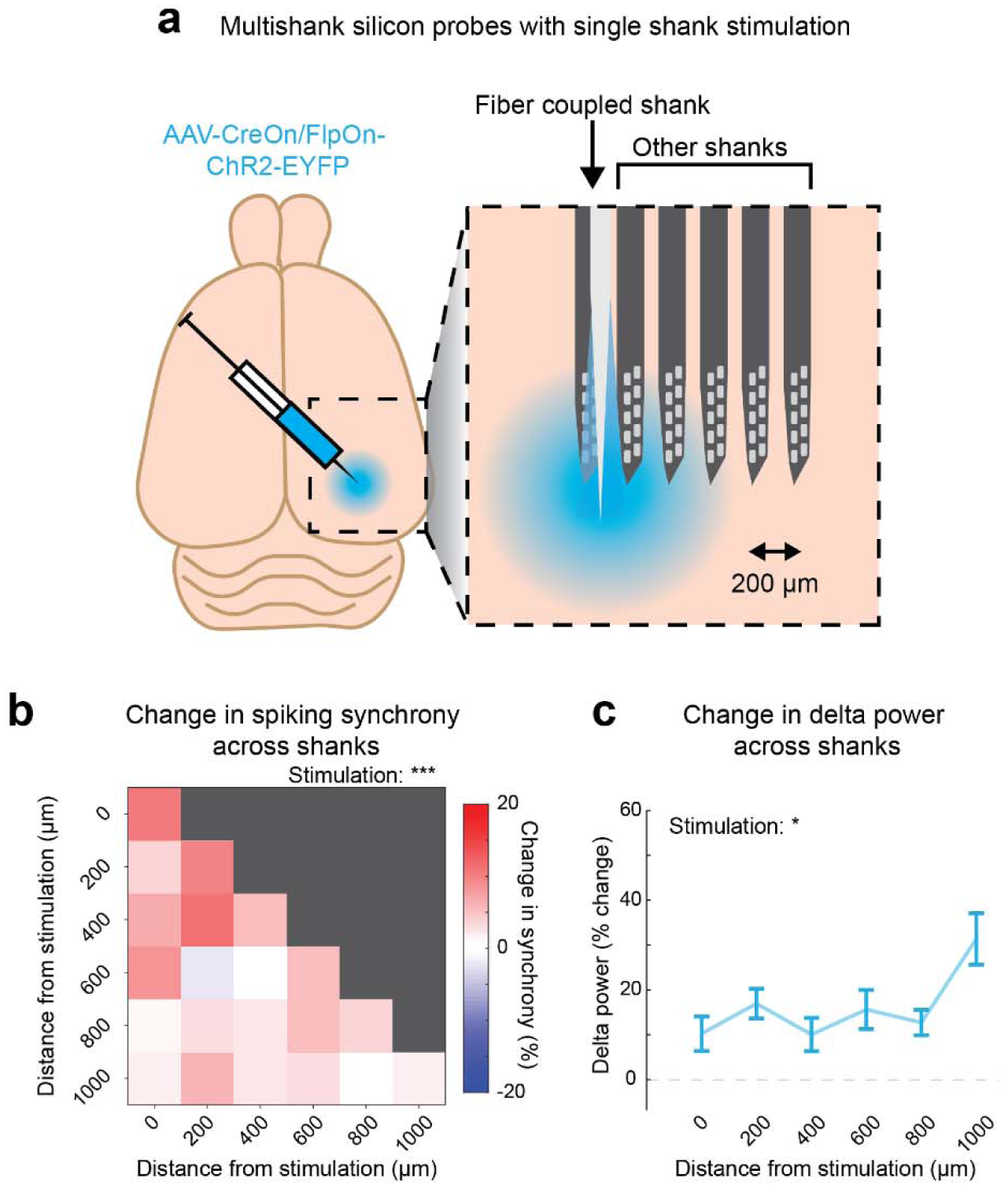
Optogenetic stimulating of Sst-Chodl cells increases synchrony across spatially distributed sites in visual cortex. **a)** Schematic of the multishank 64-channel silicon probe inserted into visual cortex of head-fixed mice, with a tapered optical attached to the most lateral (first) shank to enable single-shank optogenetic stimulation while recording units and LFPs across all shanks. **b)** Average change in cross-correlograms (CCGs) during optogenetic stimulation relative to spontaneous periods for unit pairs recorded on the stimulated shank (distance = 0) and on neighboring shanks (spaced of 200µm apart). Stimulation significantly increased spiking synchrony across the array (linear regression model: stimulation effect p < 0.001, distance effect p = 0.133), indicating that the synchrony enhancement propagated well beyond the directly stimulated site. **c)** Change in delta-band (1-4Hz) power across shanks during stimulation (linear mixed-effects model with post hoc ANOVA: stimulation effect p = 0.049, distance-stimulation interaction p = 0.270). Delta power increase across all shanks, showing that Sst-Chodl cell activation induces distributed low-frequency synchronization across visual cortex. N = 18 sessions from 8 mice. ***: p < 0.001, *: p < 0.05. Data are means; bars indicate s.e.m.

**Supplemental Figure 11.**
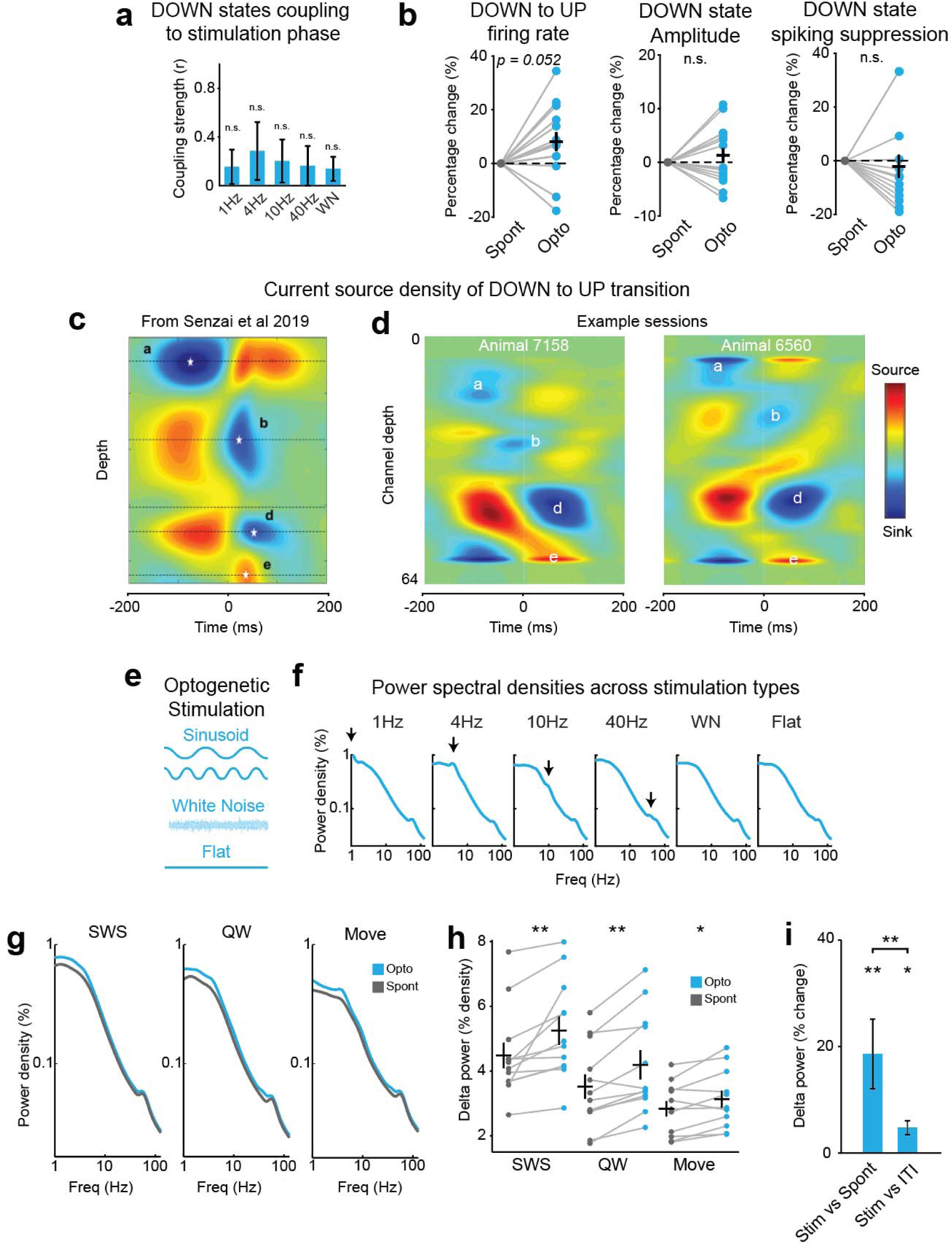
Optogenetic activation of Sst-Chodl cells enhances delta power and modulates slow-wave dynamics across behavioral states. **a)** Coupling strength of DOWN states to the phase of optogenetic stimulation (Rayleigh test for non-uniformity, n.s., p > 0.05). DOWN states did not become phase-locked to the stimulation cycle, indicating that stimulation increased synchrony without imposing a fixed phase structure. **b)** Left: change in firing rate during DOWN to UP states; middle: change in DOWN-state LFP amplitude; right: suppression of spiking during DOWN states. None of these features differed significantly between stimulation and spontaneous periods (Firing rate p = 0.052; Amplitude p = 0.496; Suppression p = 0.361; paired t-tests), suggesting that Sst-Chodl cell activation does not strongly alter the intrinsic structure of DOWN-state events. **c)** Average current source density (CSD) patterns during freely moving natural sleep from ^102^. **d)** Example CSDs during head-fixed sleep in this study, showing sinks and sources labeled (a-e) corresponding to motifs described in ^102^. These similarities indicate that DOWN/UP dynamics in the head-fixed preparation resemble natural sleep. **e)** Schematic of stimulation regimes used to probe frequency-dependent network responses. **f)** Power spectra for each stimulation protocol; arrows indicate stimulation frequencies. Optogenetic activation produces characteristic increases in low-frequency power. **g)** Change in LFP power spectra across states (SWS, quiet wake - QW, Movement) during stimulation of Sst-Chodl cells. **h)** Delta-band (1–4 Hz) power increases across all states during stimulation (SWS p = 0.008; QW, p = 0.001; Move p = 0.027; paired t-tests) , demonstrating that Sst-Chodl cells elevate low-frequency power even when animals are awake or moving. **i)** Delta power change across different epochs when comparing stimulation, ITIs, and spontaneous periods (overall p = 0.008; Stim vs. ITI p = 0.037; Stim vs. Spont p = 0.005; paired t-tests). These analyses confirm that Sst-Chodl cell activation reliably increases delta power regardless of behavioral epoch. These experiments were performed in V1 of head-fixed mice. N = 14 sessions across 10 mice. **: p < 0.01, *: p < 0.05; ns, not significant. Data are means; shading and bars indicate s.e.m.

**Supplemental Figure 12.**
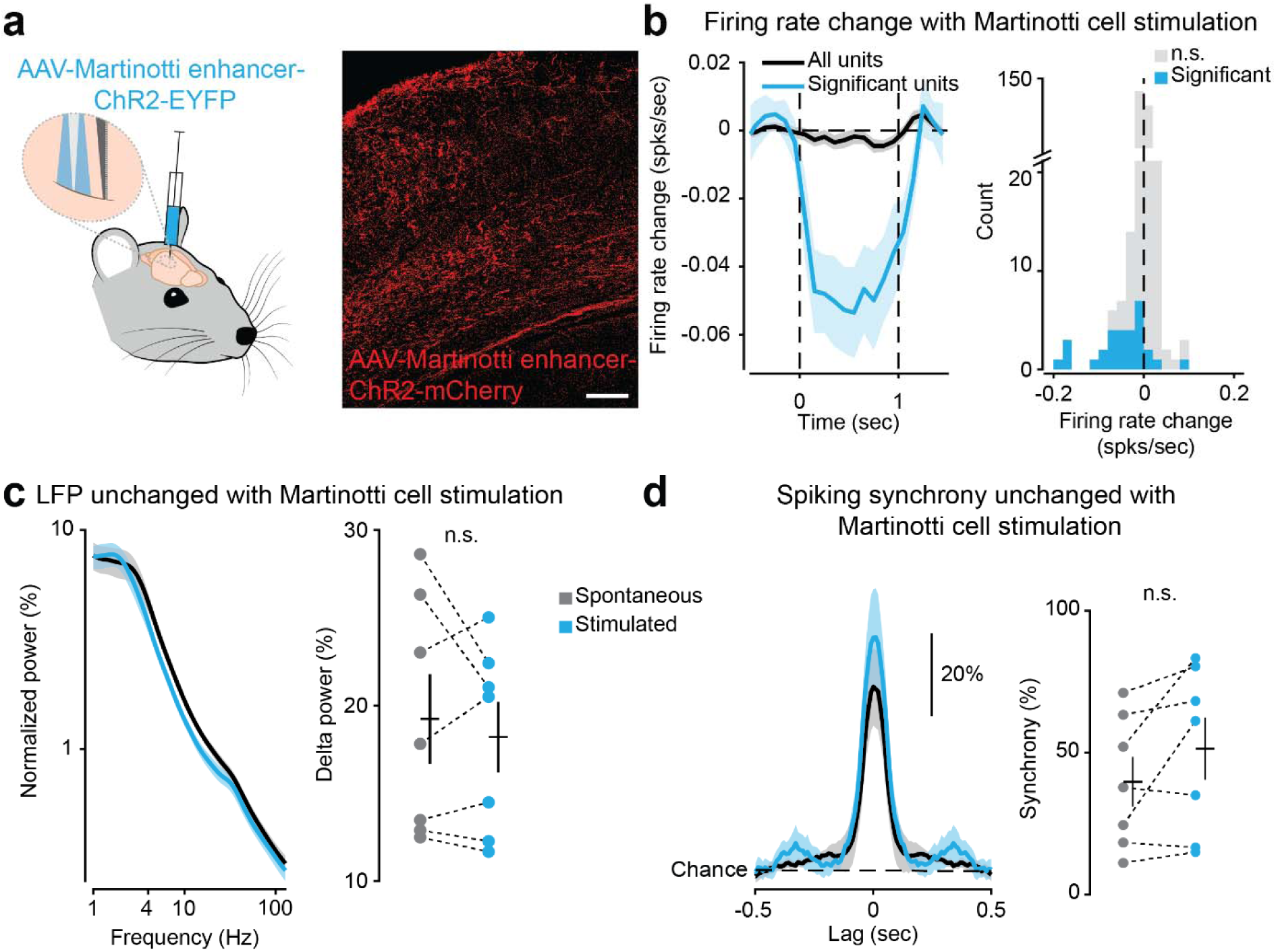
Stimulation of Martinotti (non-Chodl Sst) cells does not increase cortical synchrony. **a)** Left: schematic of the experimental strategy using an AAV carrying a Martinotti-cell–specific enhancer driving ChR2-YFP expression in V1 of head-fixed mice. Right: example ChR2 expression (scale: 100 µm). This approach selectively activates a major non-Chodl Sst subtype (Martinotti cells). **b)** Change of firing rate during Martinotti-cell stimulation (0 to 1). The firing rate of 30 out 361 units was found to be significantly changed during optogenetic activation. **c)** Left: power spectra density during spontaneous and stimulation periods. Right: quantification of delta-band (1–4 Hz) LFP power, showing no significant change with stimulation (p = 0.460, paired t-test). This indicates that Martinotti-cell activation does not increase low-frequency power. **d)** Left: average normalized cross correlogram (CCG) for unit pairs during spontaneous and stimulation epochs. Right: quantification of pairwise synchrony, showing no significant change (p = 0.100, paired t-test). Thus, unlike Sst-Chodl cells, Martinotti cells do not induce increases in spiking synchrony. N = 7 sessions across 3 mice; 361 units and 28,016 pairs. ns, not significant. Data are means; shading and bars indicate s.e.m.

**Supplemental Figure 13.**
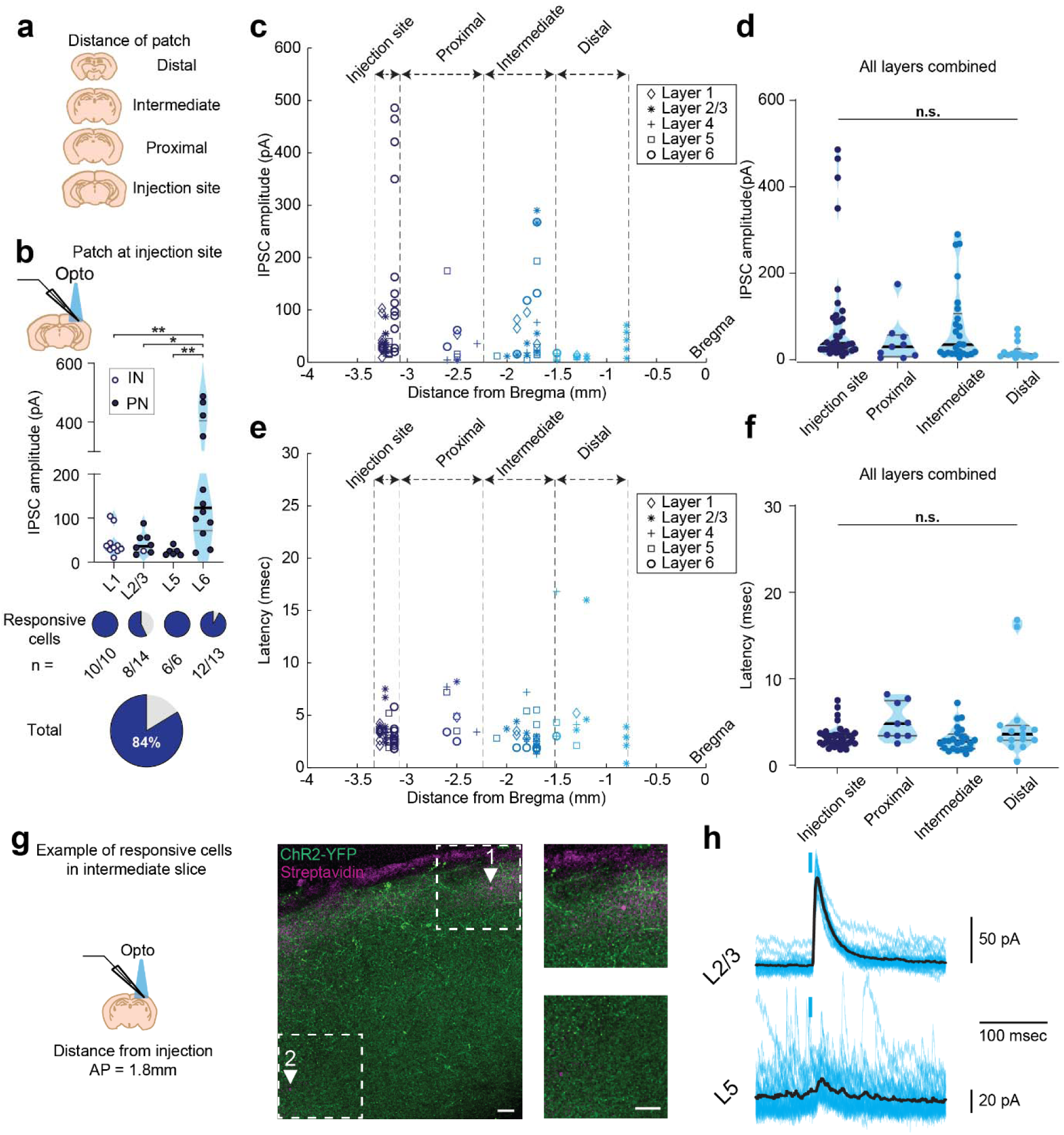
Sst-Chodl cells provide widespread monosynaptic inhibition across neocortical layers and distances. **a)** Schematic of acute slices recorded at different distances from the V1 injection site (injection site: -3.2mm from Bregma; Proximal: -3.19 to - 2.21mm; Intermediate: -2.2 to -1.51mm; Distal: -1.5 to 0.79mm). This strategy allowed assessment of long-range inhibitory connectivity of Sst-Chodl cells. **b)** Optogenetically evoked inhibitory postsynaptic current (oIPSC) recorded at the injection. Top: oIPSC amplitude across layers for putative interneurons (IN; blue/white circles) and putative pyramidal neurons (PN; black/blue circles). Bottom: pie charts showing the proportion of neurons in each layer responding to Sst-Chodl activation (One-way ANOVA interaction p = 0.001; L1 vs. L6 p = 0.005; L2/3 vs. L6 p = 0.013; L5 vs. L6 p = 0.009; other comparisons p > 0.9; n = 43 cells from N = 10 mice). Layer 1 exhibited the highest proportion of responsive cells. Layer 6 showed the largest amplitudes. These results confirm dense local inhibition across multiple layers. **c)** oIPSC amplitudes across all distances and layers. Dash lines mark boundaries between distance categories. **d)** Mean oIPSC amplitude across layers and distances (ANOVA interaction on linear mixed-effects model p = 0.794; Layer effect p < 0.001; Distance effect p = 0.795). oIPSC amplitudes varied by layer but remained detectable even at distal recording sites, consistent with widespread long-range inhibition. **e-f)** Same analyses as in (b,d) but for oIPSC latency. Latency did not differ significantly across layers or distances (ANOVA interaction p = 0.906; Layer effect p = 0.321; Distance effect p = 0.604), indicating monosynaptic connectivity throughout the sampled neocortical territory. Horizontal lines indicate median (black) and interquartile range (grey). N=16 animals, n=156 cells (Injection site, 43 cells; Proximal, 21 cells; Intermedial, 48 cells; Distal, 44 cells). **g)** Example distant slice (AP -1.8mm) showing two responsive cells filled (magenta) in L2/3 and L5, and ChR2-expressing Sst-Chodl fibers (green), demonstrating long-range innervation at distant cortical sites and fibers present in superficial layers, consistent with anatomical projections. **h)** Individual oIPSCs (blue) with averaged responses (25-30 traces; black) aligned to the light stimulation (blue rectangle) from the cell filled in (g). Strong oIPSCs from distal locations further confirm long-range monosynaptic inhibition. N = 16 animals; n = 156 cells total (Injection site: 43; Proximal: 21; Intermediate: 48; Distal: 44). ***: p < 0.001, **: p < 0.01, *: p < 0.05, ns = not significant.

**Supplemental Figure 14.**
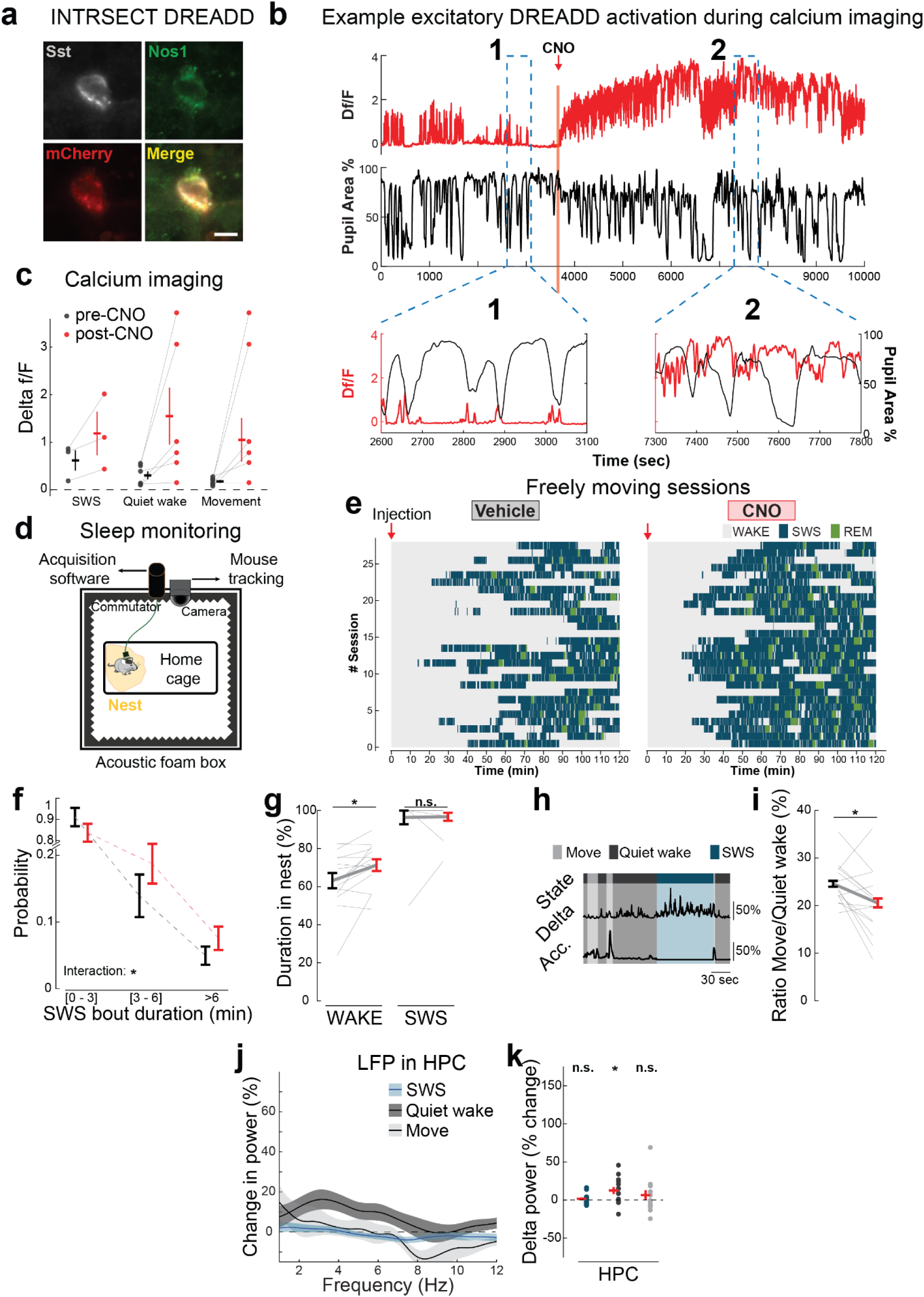
Chemogenetic activation of neocortical Sst-Chodl cells promote sleep behavior. **a)** Histological example of a neocortical Sst-Chodl cell expressing the CreOn/FlpOn excitatory DREADD (hM3Dq–mCherry). Scale bar: 20 µm. **b)** Example 2-photon recording showing increased calcium activity (red) in a DREADD-expressing Sst-Chodl cell following systemic CNO (0.5 mg/kg), along with simultaneous pupil measurements (black). CNO increased activity regardless of arousal state. **c)** Mean calcium activity across behavioral states before (pre-CNO) and 15-60 min after CNO injection (post-CNO) (n = 6 cells from 4 mice). SWS was not present in sessions from the 3 recorded cells. CNO elevated Sst-Chodl cell activity broadly across states. **d)** Schematic of sleep-monitoring paradigm for freely moving mice. **e)** Hypnogram from all sessions following Vehicle vs. CNO injections, illustrating increased SWS and REM following Sst-Chodl cell activation. **f)** Distribution of SWS bout durations following pan-neocortical activation of Sst-Chodl cells (two-way ANOVA interaction p = 0.040; 0–3 min p = 0.091; 3–6 min p = 0.118; >6 min p = 0.373). Activation shifted the distribution mainly toward more bouts. **g)** Percentage of time spent in nest during WAKE (p = 0.216) and SWS (p = 0.864), showing that nest occupancy increases during CNO treatment (red). **h)** Example of delta amplitude and accelerometer signal across Movement, Quiet Wake, and SWS. **i)** Ratio of Movement to Quiet Wake time following CNO injection (p = 0.028, paired t-test), indicating reduced wakeful activity. **j)** LFP power changes recorded in the hippocampus (HPC) during SWS, Quiet Wake and Movement. **k)** Delta band (1-4Hz) power change in HPC during SWS (blue, p = 0.376), Quiet Wake (dark grey, p = 0.016). and Movement (light grey, p = 0.341). Effects were weaker than in neocortex, suggesting preferential engagement of cortical circuits by Sst-Chodl cell activation. N = 14 mice, injections were performed during the light/inactive phase. **, p < 0.01; *, p < 0.05; ns = not significant; paired t-test for c, g, i. Data are means; shading and bars indicate s.e.m.

**Supplemental Figure 15.**
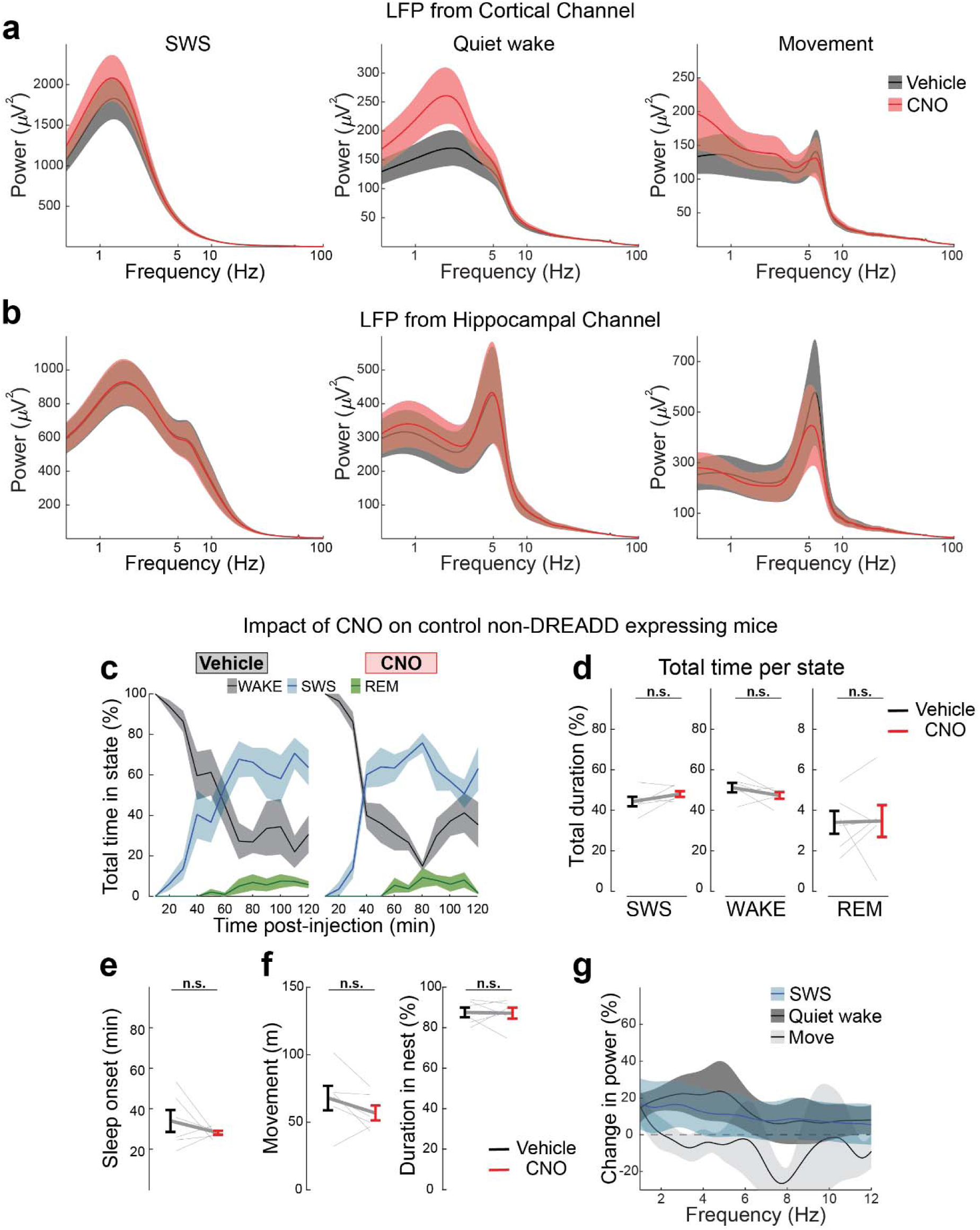
Pan-neocortical activation of Sst-Chodl cells increases cortical delta power, while CNO has no effect in control mice. **a)** Power spectral densities (PSDs) during SWS (left), Quiet Wake (center), and Movement (right) recorded from neocortical LFP channels after Vehicle or CNO injection. Chemogenetic activation of Sst-Chodl cells increased low-frequency power across behavioral states. **b**) Same analysis as in (a), but for hippocampal (HPC) channels. Delta enhancements were weaker in HPC than in neocortex, indicating preferential cortical engagement by Sst-Chodl activation. **c)** Percentage of time spent in Wake, SWS, and REM after Vehicle or CNO injection in control mice that do not express the DREADD receptor. **d)** Total time in each state during the 2-hour post-injection window for control mice (SWS p = 0.146; Wake p = 0.188; REM p = 0.931; paired t-tests). CNO alone did not alter sleep architecture in controls. **e)** Latency to sleep onset in control mice (p = 0.344), showing no effect of CNO in the absence of DREADD receptors. **f)** Total distance travelled (p = 0.145) and time spent in nest (p = 0.909) over 2 hours in control mice. No behavioral changes occurred with CNO alone. **g)** LFP power changes in control mice from neocortical channels during SWS, Quiet Wake and Movement. No significant effects were observed, confirming that the sleep-promoting and synchronizing effects require activation of Sst-Chodl cells. Ns = not significant; Paired t-test used for d-f. N = 14 mice for analyses in (a,b); N = 6 mice for analyses in (c-g). All injections were performed during the light/inactive phase. Data are means; shading and bars indicate s.e.m.

**Supplemental Figure 16.**
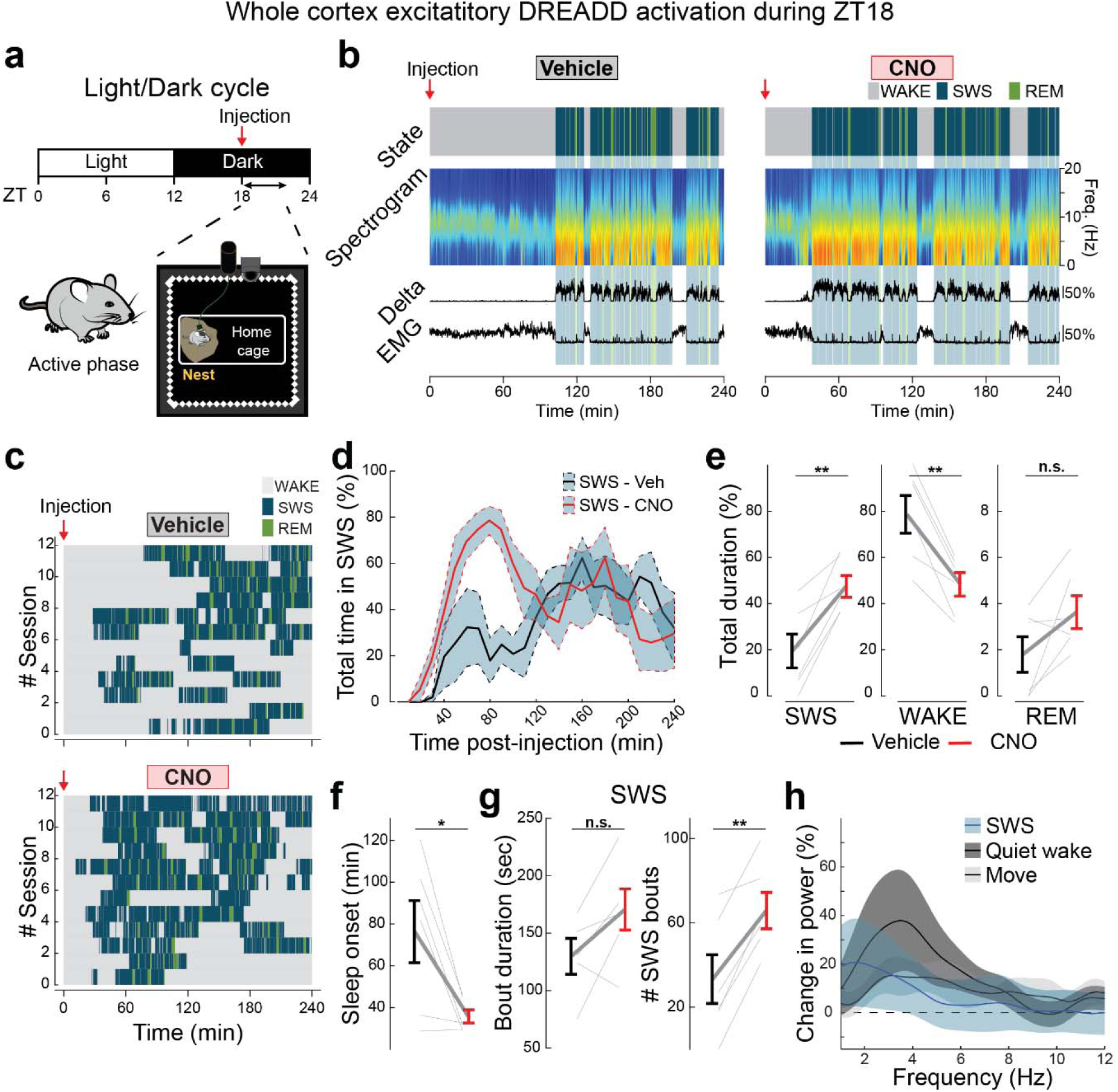
Chemogenetic activation of neocortical Sst-Chodl cells promotes sleep during the dark/active phase. **a)** Experimental design for sleep monitoring during the dark/active phase, with systemic injections performed at Zeitgeber Time 18 (ZT18), when mice are normally awake. **b)** Example recordings from one mouse during the dark/active phase after Vehicle (left) or CNO (0.5mg/kg; right), showing sleep/wake scoring and associated physiological measurements. **c)** Hypnogram for all freely moving dark/active phase sessions following Vehicle and CNO injections, illustrating a shift toward increased sleep during Sst-Chodl cell activation. **d)** Percentage of time spent in SWS after Vehicle vs. CNO. **e)** Total time in each brain state during the 2-hour post-injection window (SWS p = 0.003; WAKE p = 0.002; REM p = 0.077). CNO significantly increases SWS and reduces WAKE, with a trend toward increased REM. **f)** Latency to sleep onset (p = 0.025), showing faster transition into sleep when Sst-Chodl cells are activated. **g)** SWS bout duration (ns, p = 0.110) and numbers of SWS bouts (p = 0.002), showing that Sst-Chodl cell activation promotes sleep primarily by increasing bout number and reducing latency, rather than lengthening individual bouts. **h)** LFP power changes recorded in neocortical channels during SWS, Quiet Wake, and Movement following CNO. Activation of Sst-Chodl cells increased low-frequency power across states. Together, these results are consistent with effects observed during the light/inactive phase. **, p < 0.01; *, p < 0.05; ns = not significant. Paired t-test for all comparisons. N = 6 mice. Data are means; shading and bars indicate s.e.m.

**Supplemental Table 1.** Single-cell projection density of Sst-Chodl neurons in V1 and higher visual areas (HVAs). Summary of axonal projection patterns for individually reconstructed Sst-Chodl cells. For each neuron, the table reports soma location and the total axonal length contained in each neocortical area. All values are from the hemisphere ipsilateral to the soma, as no contralateral projections were detected. This table provides quantitative support for the broad, intracortical, long-range arborization of Sst-Chodl neurons. See excel file named **Supplemental Table 1**.

**Supplemental Table 2.** Branching and terminal projection densities of Sst-Chodl neurons. For each reconstructed Sst-Chodl cell, this table reports the neocortical areas that contain both axonal branch points and terminal endings, following conventions from prior anatomical studies^104^. Values include soma location and the total axonal length within each such area. All data are from the hemisphere ipsilateral to the soma, as no contralateral projections were observed. This table highlights the specific regions where Sst-Chodl axons form local arbor complexity and terminate. See excel file named

**Supplemental Table 3:** Complete list of area projection density map of Sst-Chodl neurons across neocortical and hippocampal areas. Table reporting the projection density (axon path length per unit volume) of reconstructed Sst-Chodl neurons across all neocortical regions defined in the Allen Common Coordinate framework, as well as hippocampal formation areas in which projections were detected. This comprehensive dataset provides a full quantitative overview of Sst-Chodl long-range intracortical innervation patterns.

| Area Name | Mean density | Density std. dev. |
| --- | --- | --- |
| rostrolateral area | 353.817744 | 86.8542906 |
| primary visual area | 254.230876 | 65.5825874 |
| anterolateral visual area | 190.110085 | 132.862149 |
| posteromedial visual area | 161.073969 | 106.692083 |
| lateral visual area | 101.274409 | 77.2172299 |
| anteromedial visual area | 100.422502 | 84.620492 |
| anterior area | 97.7034925 | 81.273239 |
| retrosplenial area, lateral agranular part | 49.2437828 | 98.0013612 |
| laterointermediate area | 28.0897375 | 39.5950979 |
| primary somatosensory area, barrel field | 27.8738788 | 16.9356931 |
| primary somatosensory area, trunk | 25.4189775 | 24.0517205 |
| posterior auditory area | 15.8636398 | 14.0018957 |
| posterolateral visual area | 14.947232 | 15.1223005 |
| primary somatosensory area, lower limb | 5.23046765 | 3.22956478 |
| primary auditory area | 5.19198356 | 7.34538855 |
| temporal association areas | 4.7131325 | 3.53264572 |
| retrosplenial area, dorsal part | 4.63443433 | 3.403158 |
| dorsal auditory area | 4.55820316 | 5.11929393 |
| retrosplenial area, ventral part | 3.61808834 | 3.83703651 |
| primary somatosensory area, unassigned | 3.48660635 | 3.12185105 |
| anterior cingulate area, dorsal part | 3.39150012 | 4.90557255 |
| postrhinal area | 3.3136601 | 3.48534805 |
| supplemental somatosensory area | 2.85778752 | 3.15858147 |
| primary somatosensory area, upper limb | 2.8159213 | 2.811427 |
| secondary motor area | 1.54072935 | 1.4655554 |
| primary somatosensory area, nose | 1.16144038 | 1.72629817 |
| primary motor area | 0.99224953 | 0.94206136 |
| ectorhinal area | 0.70558025 | 1.24361704 |
| visceral area | 0.4692631 | 0.69785985 |
| ventral auditory area | 0.39940397 | 0.25300502 |
| anterior cingulate area, ventral part | 0.31308763 | 0.36811857 |
| perirhinal area | 0.22011255 | 0.53916344 |
| primary somatosensory area, mouth | 0.15346161 | 0.23937919 |
| prelimbic area | 0.05074395 | 0.12429677 |
| agranular insular area, dorsal part | 0 | 0 |
| agranular insular area, posterior part | 0 | 0 |
| agranular insular area, ventral part | 0 | 0 |
| frontal pole | 0 | 0 |
| gustatory areas | 0 | 0 |
| infralimbic area | 0 | 0 |
| orbital area, lateral part | 0 | 0 |
| orbital area, medial part | 0 | 0 |
| orbital area, ventrolateral part | 0 | 0 |

**Supplemental Table 4:** Presynaptic partner counts of Sst-Chodl neurons across brain areas. Table listing the number of rabies-labeled presynaptic neurons were found, detected in each brain region, annotated according to the Allen Common Coordinate Framework. Counts include all areas in which retrogradely labeled cells were found, providing a comprehensive map of the upstream inputs to Sst-Chodl neurons.

| Area | Mean count | SD | Number of observations |
| --- | --- | --- | --- |
| Ammon's horn | 1 | 0 | 1 |
| *Anterior area/ layer 5* | 5.23 | 4.24 | 3 |
| Anterolateral visual area/ layer 2/3 | 12 | 0 | 1 |
| Anterolateral visual area/ layer 4 | 6 | 4.25 | 4 |
| Anterolateral visual area/ layer 5 | 5 | 4.24 | 4 |
| Anterolateral visual area/ layer 6a | 2 | 0 | 2 |
| Anteromedial visual area/ layer 1 | 1 | 0 | 1 |
| Anteromedial visual area/ layer 2/3 | 2 | 3 | 2 |
| Anteromedial visual area/ layer 5 | 1 | 0 | 1 |
| Area prostriata | 1 | 0 | 1 |
| *corpus callosum/ splenium* | 1 | 0 | 1 |
| Dorsal part of the lateral geniculate complex | 3 | 0.71 | 3 |
| Dorsal part of the lateral geniculate complex/<br>shell | 2.2 | 1.25 | 3 |
| Entorhinal area | 2 | 0 | 1 |
| Entorhinal area/ medial part/ dorsal zone | 2 | 0 | 1 |
| *Entorhinal area/ medial part/ dorsal zone/ layer 3* | 3 | 0 | 1 |
| Entorhinal area/ medial part/ dorsal zone/ layer 5 | 6 | 2.26 | 3 |
| Geniculate group/ dorsal thalamus | 2.5 | 0.71 | 4 |
| Inferior colliculus/ external nucleus | 1 | 0 | 1 |
| Lateral group of the dorsal thalamus | 5 | 1.99 | 3 |
| Lateral posterior nucleus of the thalamus | 6 | 4.55 | 4 |
| *Lateral visual area/ layer 2/3* | 2 | 1.41 | 3 |
| Lateral visual area/ layer 4 | 2 | 1.41 | 3 |
| Lateral visual area/ layer 5 | 1.3 | 0.58 | 3 |
| Lateral visual area/ layer 6a | 1 | 0 | 1 |
| Medial geniculate complex | 2 | 0 | 1 |
| Medial geniculate complex/ ventral part | 2 | 0 | 1 |
| optic radiation | 2 | 1.05 | 3 |
| *Posterior auditory area/ layer 5* | 1 | 0 | 2 |
| Posterior complex of the thalamus | 2 | 0 | 1 |
| Posterior parietal association areas | 10.5 | 6.27 | 4 |
| Posterolateral visual area/ layer 4 | 1 | 0 | 1 |
| Posterolateral visual area/ layer 5 | 1 | 0 | 1 |
| posteromedial visual area/ layer 1 | 3 | 1.25 | 3 |
| posteromedial visual area/ layer 2/3 | 1 | 0 | 1 |
| *posteromedial visual area/ layer 5* | 2 | 1.41 | 3 |
| Postrhinal area/ layer 5 | 2 | 0 | 1 |
| Primary auditory area/ layer 2/3 | 1 | 0 | 1 |
| Primary auditory area/ layer 4 | 1 | 0 | 1 |
| Primary somatosensory area/ barrel field/ layer 2/3 | 7.3 | 4.62 | 4 |
| *Primary somatosensory area/ barrel field/ layer 4* | 3 | 2.83 | 4 |
| Primary somatosensory area/ barrel field/ layer 5 | 3.5 | 1.91 | 4 |
| Primary somatosensory area/ barrel field/ layer 6b | 1 | 0 | 1 |
| Primary visual area/ layer 1 | 4.6 | 1.41 | 3 |
| Primary visual area/ layer 2/3 | 56.5 | 26.98 | 4 |
| *Primary visual* area/ layer 4 | 21.3 | 10.34 | 4 |
| Primary visual area/ layer 5 | 31.5 | 14.18 | 4 |
| Primary visual area/ layer 6a | 17.5 | 7.78 | 4 |
| Primary visual area/ layer 6b | 4 | 2.26 | 4 |
| Retrosplenial area/ dorsal part/ layer 5 | 1 | 0 | 1 |
| Retrosplenial area/ lateral agranular part/ layer 5 | 1 | 0 | 1 |
| *Retrosplenial* area/ lateral agranular part/ layer 6a | 1 | 0 | 1 |
| Retrosplenial area/ ventral part layer 3 | 1 | 0 | 1 |
| Retrosplenial area/ ventral part/ layer 5 | 1 | 0 | 1 |
| Rostrolateral area/ layer 2/3 | 29 | 12.12 | 4 |
| Rostrolateral area/ layer 4 | 11.7 | 0.58 | 4 |
| Rostrolateral area/ layer 5 | 8.3 | 4.27 | 4 |
| Rostrolateral area/ layer 6a | 9.5 | 0.71 | 3 |
| Rostrolateral area/ layer 6b | 1 | 0 | 1 |
| Somatosensory areas | 7.3 | 2.12 | 3 |
| Subiculum | 1 | 0 | 1 |
| Supplemental somatosensory area/ layer 2/3 | 3 | 2.2 | 3 |
| Supplemental somatosensory area/ layer 4 | 3 | 0 | 1 |
| Supplemental somatosensory area/ layer 5 | 1 | 0 | 1 |
| supra-callosal cerebral white matter | 1 | 0 | 1 |
| Temporal association areas/ layer 5 | 1 | 0 | 1 |
| Thalamus/ polymodal association cortex related | 2.1 | 0.96 | 4 |
| Thalamus/ sensory-motor cortex related | 2.2 | 0.71 | 4 |

**Supplemental Video 1: Whole-brain visualization of Sst-Chodl cells labeled in primary somatosensory cortex (S1).** Three-dimensional rendering of Sst-Chodl cells labeled by intersectional oScarlet AAV injected into S1 and imaged in a whole-mount cleared brain. The video illustrates the dense local arborization and long-range intracortical projections of Sst-Chodl cells across neocortical regions.

**Supplemental Video 2: Imaging of a DREADD-expressing Sst-Chodl cell prior to CNO administration.** Simultaneous facial videography and 2-photon calcium imaging of a neocortical Sst-Chodl cell expressing excitatory DREADD. Pupil diameter (yellow) and calcium activity (red) are shown along with a real-time indicator (blue line). During periods of reduced arousal, reflected by pupil constriction, the Sst-Chodl cell exhibits increased activity, illustrating the natural state dependence of these neurons before chemogenetic activation.

**Supplemental Video 3: Chemogenetic activation disrupts the normal arousal–activity relationship in Sst-Chodl cells.** Simultaneous facial videography and 2-photon imaging of a neocortical Sst-Chodl cell expressing excitatory DREADD after systemic CNO administration. Pupil diameter (yellow) and calcium activity (red) are displayed along with a real-time indicator (blue line). Following CNO injection, the cell becomes persistently active, and the typical coupling between low arousal (pupil constriction) and increased Sst-Chodl cell activity is abolished, demonstrating effective chemogenetic activation.

